# Single-cell mosaicism analysis reveals cell-type-specific somatic mutational burden in Alzheimer’s Dementia

**DOI:** 10.1101/2022.04.21.489103

**Authors:** Maria Kousi, Carles Boix, Yongjin P. Park, Hansruedi Mathys, Samuel Sledzieski, Zhuyu Peng, David A. Bennett, Li-Huei Tsai, Manolis Kellis

**Affiliations:** Computer Science and Artificial Intelligence Laboratory, Massachusetts Institute of Technology, Cambridge, MA, USA; Broad Institute of MIT and Harvard, Cambridge, MA, USA; Computational and Systems Biology Program, Massachusetts Institute of Technology, Cambridge, MA, USA; Picower Institute for Learning and Memory, Cambridge, MA, USA; Department of Brain and Cognitive Sciences, Cambridge, MA, USA; Rush Alzheimer’s Disease Center, Rush University Medical Center, Chicago, IL, USA

**Author notes:** Equal contributors (Maria Kousi, Carles Boix).

## Abstract

Despite significant advances in identifying genetic drivers of neurodegenerative disorders, the majority of affected individuals lack molecular genetic diagnosis, with somatic mutations proposed as one potential contributor to increased risk. Here, we report the first cell-type-specific map of somatic mosaicism in Alzheimer’s Dementia (AlzD), using 4,014 cells from prefrontal cortex samples of 19 AlzD and 17 non-AlzD individuals. We integrate full-transcript single-nucleus RNA-seq (SMART-Seq) with matched individual-level whole-genome sequencing to jointly infer mutational events and the cell-type in which they occurred. AlzD individuals show increased mutational burden, localized in excitatory neurons, oligodendrocytes, astrocytes and disease-associated “senescent” cells. High-mutational-burden cells showed mutational enrichment and similar single-cell expression profiles in AlzD cases versus non-AlzD individuals, indicating cellular-level genotype-to-phenotype correlation. Somatic mutations are specifically enriched for known AlzD genes, and implicate biologically meaningful cell-type specific processes, including: neuronal energy regulation, endocytic trafficking (*NEFM*), lipid metabolism (*CNP, CRYAB*), proteostasis (*USP34*), cytoskeleton, and microtubule dynamics (*MACF1*).

## Introduction

Late-onset neurodegenerative disorders, including Alzheimer’s Dementia (AlzD), Parkinson’s disease, Huntington’s disease, and Amyotrophic Lateral Sclerosis are primarily sporadic, arising through the combined effect of multiple environmental risk factors and many weak-effect common alleles^1,2^, though heritable forms have also been described, stemming from a small number of strong-effect rare alleles^3–6^. Hindering the search for effective therapeutic interventions, our current knowledge of genetic contributors is still greatly incomplete, in part due to their complex etiology, heterogeneous level of affectedness, diverse symptoms, and the interplay between genetic and environmental factors^7,8^.

Several genetic contributors to AlzD have been elucidated, including: the *E4* haplotype of the *ApoE* gene locus^9^; at least 25 loci from genome-wide association studies (GWAS)^1,2^, including *BIN1*^*10*^, *PICALM*^*11*^, *CLU*^*11,12*^, *CR1*^*12*^; a small number of genes discovered from family pedigrees, involving *APP, PSEN1, PSEN2*^*3–6*^; and an increasingly-large set of coding and non-coding loci from ongoing exome-sequencing and whole-genome sequencing (WGS) studies^13,14^. With the exception of the ApoE locus, genetic results in AlzD follow the commonly-observed inverse relationship between allele frequency and effect size, in the continuum between high-frequency weak-effect variants (from GWAS) and low-frequency strong-effect variants (from family studies), due to the impact of selection, maintaining strong-effect variants at low frequencies. Despite these major advances however, the vast majority of AlzD cases lack any detectable genetic basis.

Beyond inherited genetic variants, genetic contributors to disease can include somatic mutations incurred either during early embryonic development^15^ (thus affecting a large fraction of cells and tissues), in later stages of embryogenesis (thus confined to a specific tissue or cell type), or during childhood, adulthood, and late-life (thus affecting only a small number of cells). These mutations can stem from environmental exposures, disease-associated DNA damage, disruption of DNA repair processes, or through the natural process of error-prone mitotic cell divisions. The phenomenon describing such somatic mutations is referred to as mosaicism, given that the affected individual is a mosaic of cells of different genetic makeup, including both risk and protective alleles of disease loci. Unlike inherited mutations, late-stage somatic mutations are not subject to the strong selective pressures acting on germline variation, and thus can result in strong gene-perturbing effects, which can have a disproportionate impact on cellular phenotype. Increased somatic mutational burden in individual cells can increase the chance of single-cell dysregulation. Increasing numbers of dysregulated cells can subsequently manifest as tissue- and organ-level disruptions, providing an additional path for disease onset and progression, beyond inherited genetic components and environmental contributors.

Though significant progress has been made in studying the relevance of mosaicism in disease, with prominent examples from cancer and clonal hematopoietic malignancies^16–18^, congenital heart disorders^19^, and skin disorders^20^, the inaccessibility of brain tissue has hindered the study of mosaicism in neurodevelopmental and neurodegenerative disease^21,22^. Bulk DNA sequencing has been used to recognize high-frequency somatic mutations in autism spectrum disorders^23–25^ and in hemimegalencephaly^26^, from their partial representation in WGS reads. For DNA-damage-repair disorders, a single-cell DNA sequencing study showed increased mutations in nine cases vs. controls^27^, profiling three to six neurons from each affected individual. Thus, it remains unclear whether somatic mosaicism plays a role in neurodegeneration more broadly (outside DNA-damage-repair disorders), whether increased mutations are also found in non-neuronal cells (which are increasingly recognized to play important roles in AlzD^28^), and how DNA perturbations relate to gene expression changes at the single-cell level, and ultimately to phenotypic effects in brain function and cognition. Deep whole-genome sequencing in individual sorted neuronal cells from AlzD and control individuals has shown an increase in somatic mutational burden^29^, but the transcriptional state context of these cells remains unknown, making it impossible to correlate transcriptional phenotypic changes associated with somatic burden in individual cells.

Here, we report the first single-cell joint map of somatic mutation and gene expression patterns in AlzD, using 4,014 cells from prefrontal cortex samples of 36 human *post mortem* brains with matched WGS data, including 19 AlzD and 17 non-AlzD individuals based on cognitive clinical diagnosis. We profile full-transcript single-nucleus RNA sequencing with SMART-Seq, and develop a new method to jointly infer somatic mutations and single-cell gene expression levels, enabling us to map mosaic mutations onto specific cell types and cellular states in which they occur. We find that AlzD individuals show higher overall mutational burden, higher damaging-variant burden for missense and stop-codon mutations, and higher cell-type-specific burden for excitatory neurons, oligodendrocytes, astrocytes, and a newly-described class of “senescent” cells. Somatic mutations are enriched for known AlzD genes, and for specific novel and biologically-meaningful genes and pathways, including: neuronal energy regulation, endocytic trafficking, lipid metabolism, proteostasis, cytoskeleton, and microtubule dynamics. Overall, our study uncovers the role of cell-type specific mutational burden in AlzD, reveals candidate new AlzD target genes and pathways, and provides a general framework for understanding somatic variation and its phenotypic associations at the single-cell level in human disease.

## Results

### Cohort assembly and variant detection

We obtained *post-mortem* human prefrontal cortex brain samples from 47 Religious Orders Study and Rush Memory and Aging Project (ROSMAP)^30^ individuals (**Fig. 1a**), classified into 22 AlzD and 25 non-AlzD individuals, of which 36 also have whole-genome-sequencing (30x coverage)^31^.

**Figure 1.**
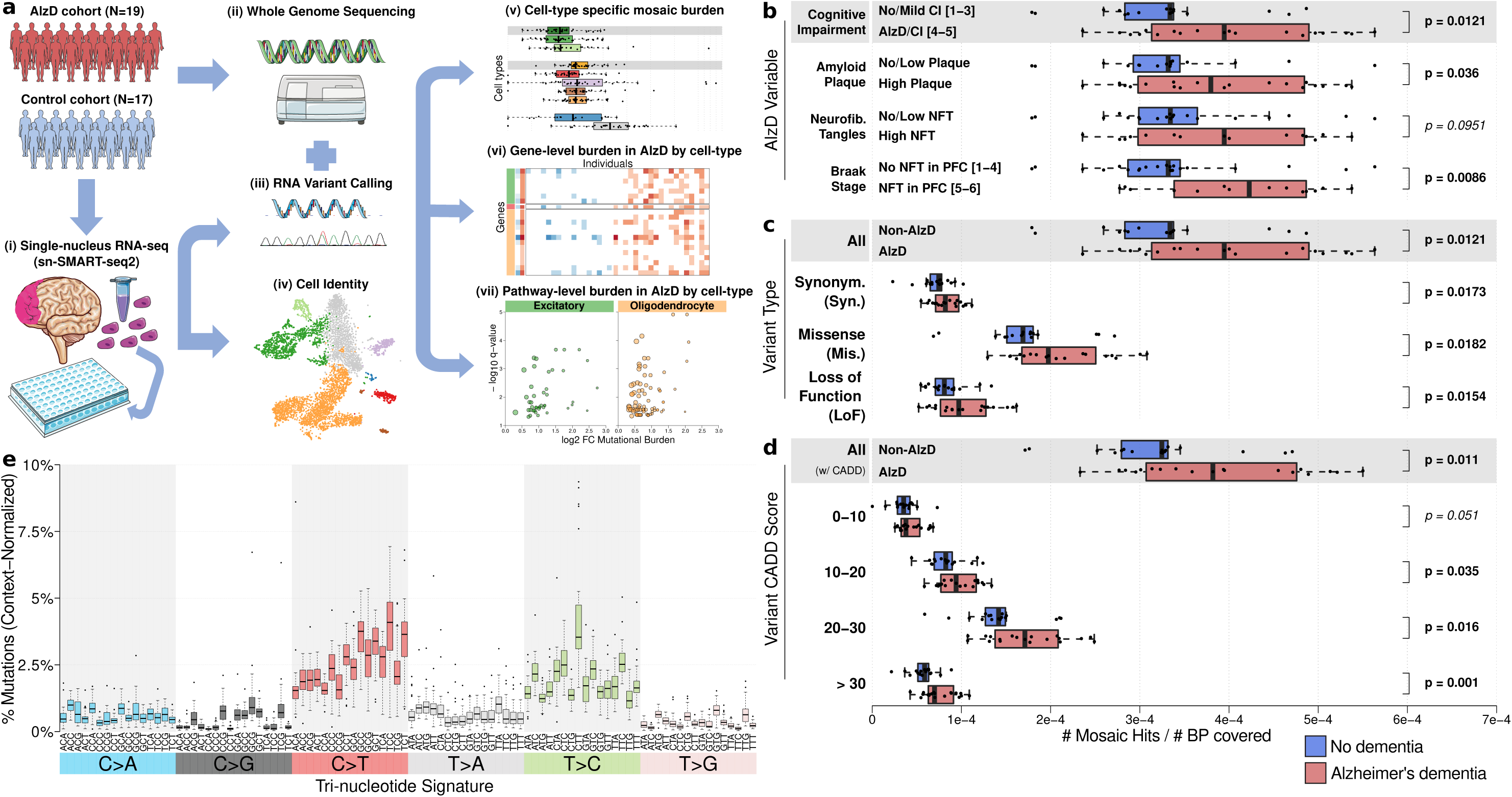
Somatic mutation burden in Alzheimer’s Dementia (AlzD). **a. Overview**. Across 36 post-mortem prefrontal cortex samples from 19 AlzD and 17 non-AlzD individuals (i), we profile full-transcript single-nucleus RNA-seq (SMART-seq2) (ii) and whole-genome sequencing (iii) to jointly infer mosaic mutations and cell identity (iv) across 4,014 cells, which we use to infer cell-type-specific mosaic burden (v), gene-level burden (vi) and pathway-level burden (vii). **b-d**. Increased mosaic mutational burden (x-axis) between AlzD (red) and non-AlzD (blue) individuals for different ascertainment variables (**b**), types of mutation (**c**), and predicted variant effect (**d**, CADD score). **e**. Normalized frequency (y-axis) of somatic mutations for different single-nucleotide variant tri-nucleotide contexts (columns) across individuals (points) highlights aging-related signatures (C-to-T followed by T-to-C).

We used well-based single-nucleus full-transcript profiles (Smart-Seq2 v2.0^32^) across all 47 individuals, and obtained 6,180 cells (5,421 after QC). We used known marker gene expression to annotate 2,121 oligodendrocytes (39.1%), 1,170 excitatory neurons (21.6%), 255 microglia (4.7%), 221 inhibitory neurons (4.1%), 220 astrocytes (4.1%), 94 OPCs (1.7%), and 40 endothelial cells (0.7%). We also identified a cluster of 1,300 cells (24%) that we refer to here as “senescent” with reduced transcription, loss of cell-type-identity markers, and expression of a subset of senescence-associated markers (AXL, SENP7), and positioned between neurons and oligodendrocytes in the two-dimensional t-SNE cell embedding. However, these did not express all classical senescence markers (e.g. CDKN1A/2A), suggesting a possibly distinct neuronal and oligodendrocyte pre-apoptotic state^33^. We only analyzed somatic mutations for the subset of 36 individuals with whole-genome sequence (encompassing 4,040 cells: 935 Excitatory neurons, 167 Inhibitory neurons, 159 Astrocytes, 188 Microglia, 74 oligodendrocyte progenitor cells (OPC), 1382 Oligodendrocytes, 32 Endocytes, 1103 Senescent cells; see Methods).

For the subset of 36 individuals with whole-genome sequencing data (4,014 cells), we inferred somatic mutations in each of their cells (GATK HaplotypeCaller, using germline-depth-adjusted thresholds, **Supplementary Fig. S1**), merged all mutations, excluded known AlzD SNPs and germline variants, and kept only exonic mutations present in >20% of reads for that cell (**see Methods**), resulting in an initial set of 66,664 mutational calls across all cells and all individuals. After excluding blacklisted regions/genes, RNA editing sites, mutations shared across >1 individuals, ExAC variants with MAF>1%, hyper-mutated cells, and merging calls within 1-3bp of another, resulting in 55,447 mutational calls (see Methods).

These mutations show high deleteriousness, indicative of relaxed selection, and consistent with their somatic occurrence in only one tissue and a subset of cells. The majority (51.3%) were amino-acid changing (missense), 24.1% loss-of-function (nonsense/stop, frameshifting, splice-disrupting), 21.5% amino-acid preserving (synonymous), and 3.1% other (inter/intragenic) (**Supplementary Fig. S2a**), with consistent frequencies across individuals and sequencing depths (**Supplementary Fig. S2b,c**). They also showed high mutation deleteriousness (CADD)^34^ scores (average=22, top 0.63% of genome), and their non-synonymous to synonymous rate^35^ deviates significantly from the baseline neutral selection (*d*N/*d*S=2.5) (**Supplementary Fig. S2e**).

### Enrichment for somatic changes in AlzD patients

Summing across all cell types, we found that somatic changes are enriched in AlzD vs. non-AlzD brains, when ascertained using cognitive impairment (Cogdx 4-5 vs. 1-3, p=0.0121; **Fig. 1b**), β-amyloid load (p=0.036), and Braak stage^36^ which measures neurofibrillary tangle (NFT) abundance across cortical areas (stage 5-6 marking NFT diffusion across neocortex vs. stage 1-4 marking NFTs restricted to entorhinal cortex and hippocampus, p=0.0086) (**Fig. 1b**). However, overall PHFtau NFT density^37^ did not show a significant difference in mutational burden (p=0.0951), highlighting the importance of clinical diagnosis and precise localization measures (see **Supplementary Fig. S3**).

We found higher somatic burden in AlzD vs. non-AlzD across all types of mutations, including synonymous (+18.1%, p=0.0173), missense (+22.6%, p=0.0182), and LoF (+25.7%, p=0.0154) (**Fig. 1c**), with the enrichment in AlzD being greater with stronger deleteriousness scores, from +16% for CADD 10-20 (p=0.035) to +22.3% for CADD 20-30 (p=0.016), to +32% for CADD>30 (p=0.001) (**Fig. 1d**). Across both AlzD and non-AlzD, changes were enriched for mutational signatures of the aging brain (C-to-T and T-to-C transitions), suggesting that burden was driven by differences in mutational abundance, rather than distinct mutational processes between AlzD and non-AlzD (**Fig. 1e, Supplementary Fig. S4a-c**)^27,29,38^.

### Cell-type identification through transcriptomic analysis

We next analyzed these mutations in the context of their cell-type-identity, leveraging the unique nature of our dataset. We annotated the cell type of our 4,040 cells across 36 whole-genome-sequenced individuals with mutational burden information, in the context of all 5,421 cells from all 47 individuals (**Fig. 2a, Supplementary Fig. S5a**). We used known marker genes to annotate seven well-established cell types, including: excitatory neurons (*CAMK2A, NRGN, SLC17A7*), inhibitory neurons (*GAD1, GAD2*), astrocytes (*AQP4, GFAP*), microglia (*C3, CD74, CSF1R)*, oligodendrocytes (*MBP, MOBP, PLP1*), OPCs (*PDGFRA, VCAN)*, and endothelial cells (*FLT1, CLDN5*) (**Fig. 2b, Supplementary Fig. S5b-d**), with each cluster showing substantial contributions by multiple individuals (**Supplementary Fig. S6,S7**). We also annotated a transcriptionally-distinct cluster of senescent cells (**Fig. 2a**), bearing resemblance to neurons and oligodendrocytes, but lacking clean cell-type identity (**Supplementary Fig. S5c-e, S8**). Senescent cells were enriched in AlzD individuals (across all ascertainment parameters, **Fig. 2c**), and also in female individuals (**Fig. 2c,e**). Oligodendrocytes were depleted in cases and in female individuals (**Fig. 2c-d**), likely reflecting higher oligodendrocyte vulnerability in AlzD, and consistent with our recently-reported AlzD-associated and sex-specific down-regulation of myelination pathways^28^.

**Figure 2.**
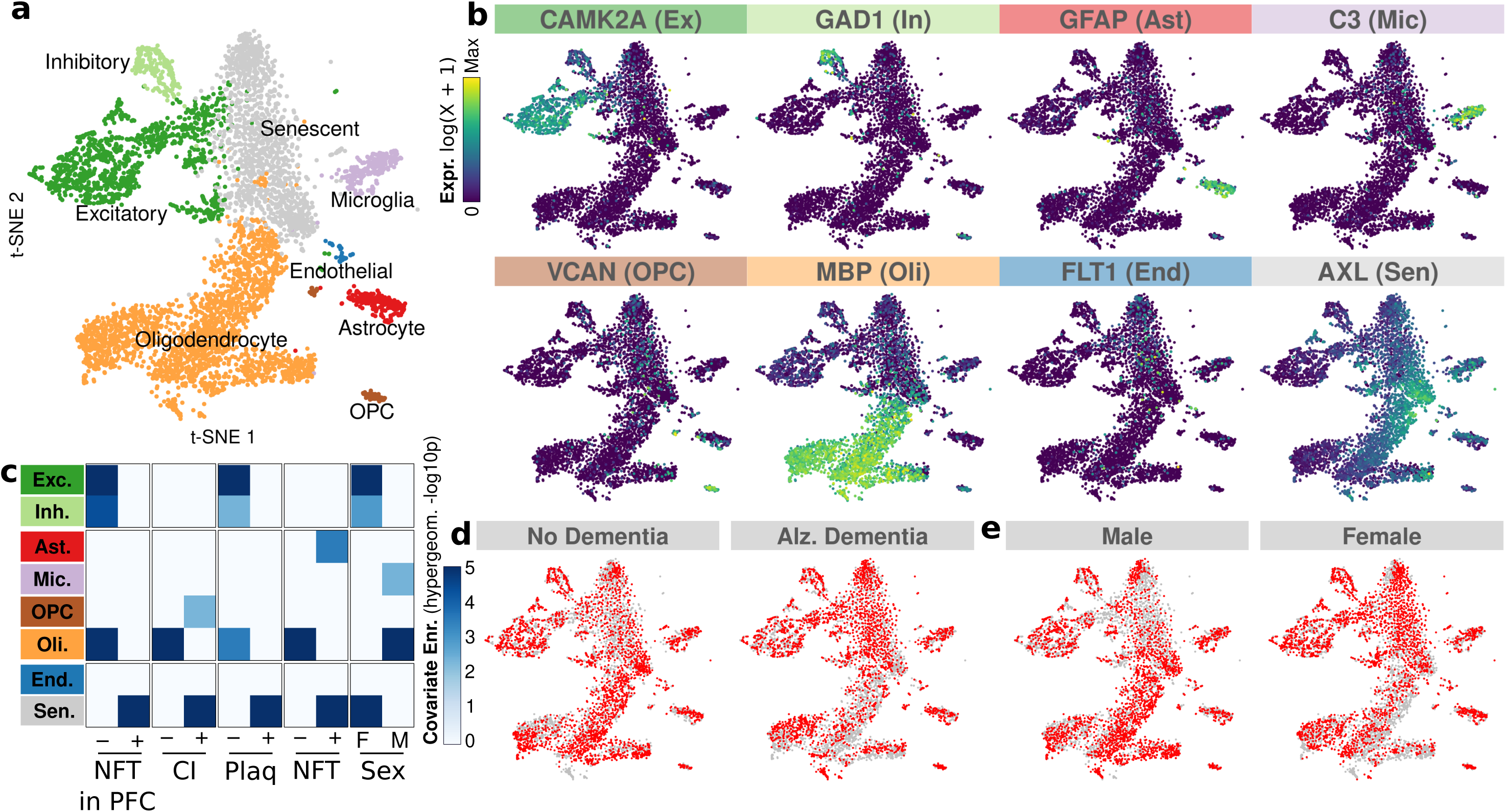
Transcriptomic landscape of Alzheimer’s dementia. **a**. Gene expression landscape (t-SNE) and cell type annotation (colors) across 5,421 cells from all 47 individuals. **b**. Representative cell-type-specific marker gene expression (color scale). **c**. Enrichment (heatmap) for each cell type (rows) across phenotypic classes (columns) and sex. **d,e**. Expression landscape (t-SNE) of cells by phenotype (d) and sex (e).

### Cell-type specific enrichment of mosaic changes in the aging brain and in AlzD

We next used these cell type annotations to evaluate the cell-type-specific mutational burden of each cell type, first focusing on overall burden irrespective of disease status. We found that glial cells show a 34.6% higher mutational burden than neuronal cells (p=3.3e-5), consistent with their continued proliferation, in contrast to post-mitotic neurons (**Fig. 3a**). Mosaic burden for both neuronal and glial cells correlated with age (r=0.09 and r=0.26) (**Fig. 3b**), consistent with the enrichment we found for aging-associated mutational signatures (**Fig. 1e**). Senescent cells harbored a 1.81-fold higher mutational load than neurons (p=2e-7) and 1.34-fold higher than glia (p=6.6e-4), suggesting that mutational load may be in part contributing to their senescent status. Male and female subjects did not show significant differences in mutational burden (**Supplementary Fig. S9**).

**Figure 3.**
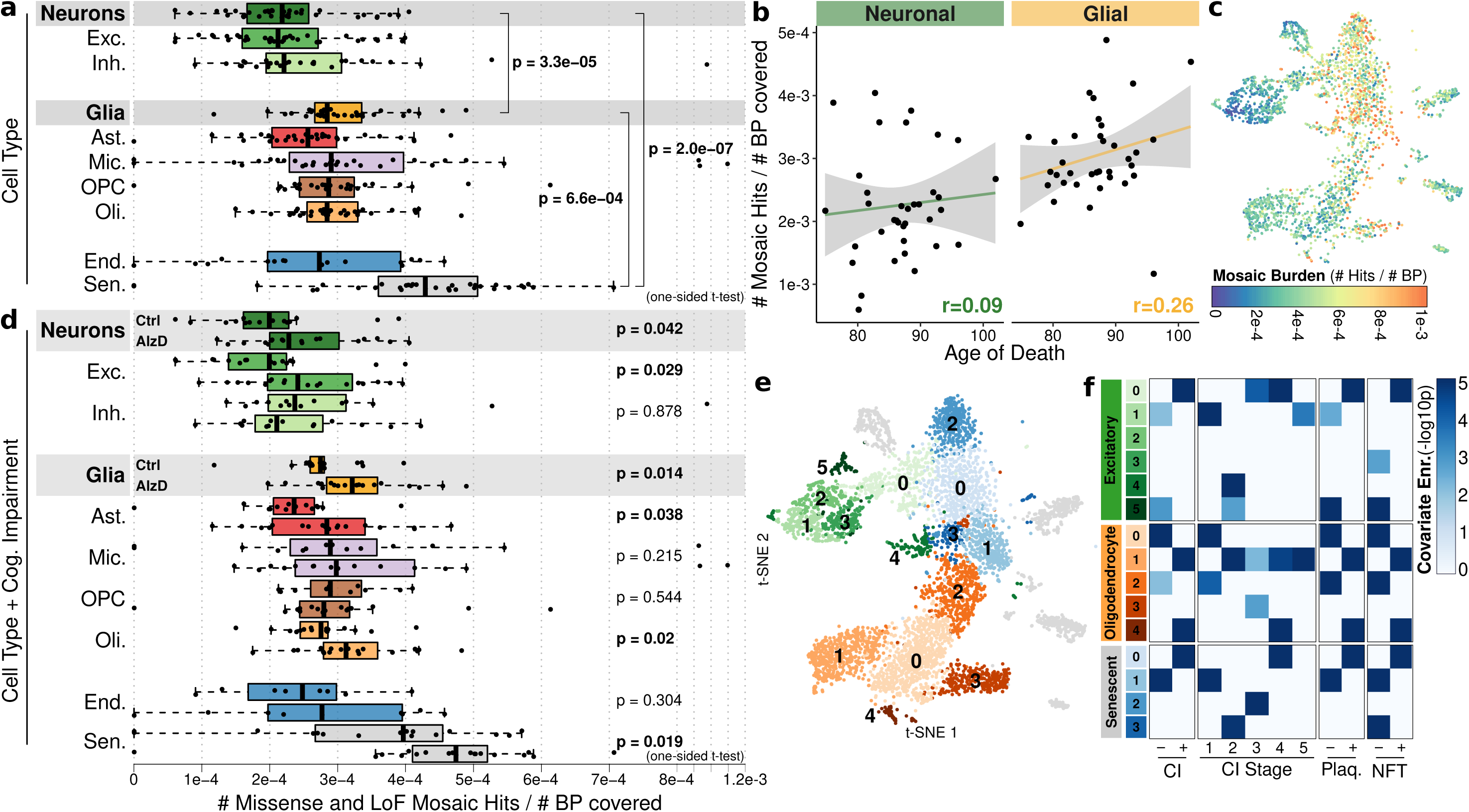
Mutational burden distribution. **a**. Mutational burden distribution (x-axis) across individuals (points) for each cell type (rows). **b**. Mosaic burden (y-axis) by age (x-axis) with linear model fit (line) and 95% confidence interval (shade). **c**. Mosaic burden (shade) across cells with mutations (points). **d**. Mosaic burden differences between Non-AlzD (top) vs. AlzD (bottom) individuals by cell type (color). **e**. Sub-clustering of most-abundant cell types. **f**. Phenotypic enrichment for cell subclusters (hypergeometric test vs. main cluster).

We next studied cell-type-specific differences in mutational burden between AlzD and non-AlzD individuals. Those with AlzD showed significantly-enriched mutational burden in neurons (24% increase, p=0.042), glia (18.5%, p=0.014), and senescent cells (34%, p=0.019) (**Fig. 3c-d**), with the neuronal signal driven primarily by excitatory neurons (30%, p=0.029), and the glial signal by astrocytes (24%, p=0.038) and oligodendrocytes (17.5%, p=0.02). These results indicate that our observed overall increase in mutational burden of AlzD individuals does not stem solely from a difference in cell type composition, but is instead reflected in each of several cell types individually. Not surprisingly, these results indicate that neurons (which have been the primary focus of previous studies of brain mosaicism^22,27^) are not the sole target of somatic mutations in AlzD, and that glial cells can show equally-strong mutational burden and disease enrichment. Our results also show that somatic variants implicate both neuronal and glial cell types, in contrast to weak-effect common variants from GWAS at the other extreme of the allele-frequency spectrum, which enrich exclusively in microglial enhancers^39,40^.

### Increased mutational burden correlates with gene expression changes at the single-cell level

For all major cell types, greater mutational burden at the single-cell level correlated with altered single-cell expression patterns (**Fig. 3e-f, Supp. Fig. S11**), indicating a relationship between genetic perturbations (somatic mutations) and phenotypic outcome (gene expression) at the cellular level. Unbiased expression-based sub-clustering (**Supplementary Fig. S10**) of excitatory neurons, oligodendrocytes, and senescent cells without any guidance from mutational information (**Fig. 3f**), resulted in cell states with significant differences in burden (**Supplementary Fig. S11a)**.

Moreover, greater mutational burden correlated with both greater enrichment in AlzD vs. non-AlzD individuals (**Supplementary Fig. S11b**), and increased gene-expression proximity to senescent cells (**Fig. 3e**), indicative of a gradient of mutational burden across cell-state space (**Fig. 3c**). For example, excitatory neuron subcluster Exc0 vs. Exc1 showed significantly-higher mutational burden (P=3.6e-9), significantly-higher AlzD vs. non-AlzD enrichment (P=2.98e-7), and highest proximity in gene-expression space to senescent cells (**Fig. 3e**). Similarly, Sen0 vs. Sen1 showed higher burden (P=0.026), higher AlzD vs. non-AlzD enrichment (p=1.13e-7), and gene expression patterns more distant from other cell types. For oligodendrocytes, Oli1 showed higher burden and higher AlzD vs. non-AlzD enrichment, though its expression patterns were shifted away from the senescent cluster, possibly indicative of disease response as we previously reported^28^.

Beyond these cluster-level correlations between greater burden and gene expression changes, we found a cell-level correlation between the two, with a gradient of mutational burden along the expression-based two-dimensional manifold t-SNE clustering of cells (**Fig. 3c**), with mutational burden highest for senescent cells and progressing to lower burden in the periphery of the t-SNE plot. Even within each subtype, the previously-noted greater burden in AlzD individuals continued to hold (**Supp Fig. S11c**), but was only significant for the largest cell states due to the decrease in power.

### AlzD-specific mosaic mutational burden concentrates in AlzD-related genes and pathways

In addition to showing higher overall mutational burden, AlzD individuals showed a strong concentration of somatic mutations in specific genes and biological processes, which was not seen for normotypic individuals, providing insights on how they contribute to AlzD pathology in individuals carrying these mutations. In total, 21 genes showed gene-level increased mutational burden in AlzD (**Fig. 4a, Supplementary Figure S12**; **Supplementary Table S1**). In excitatory neurons, seven genes showed greater mutational burden in AlzD, with roles in: protein degradation (*USP34, MYCBP2*), DNA damage (*SETX*)^41^, axon growth and transport (*NEFM*^*42*^), endoplasmic reticulum tubule turnover (*CCPG1*^*43*^), and synaptic function (*CYFIP2*^*44*^, *PCDH9*^*45,46*^). In oligodendrocytes, 13 genes with greater mutational burden in AlzD include: myelin-associated genes (phosphodiesterase *CNP*^*47*^, chaperone *CRYAB*^*48*^); dendrite development (*DOCK9*^*49*^); cytoskeleton/scaffold components (axonal growth gene *FEZ1*^*50*^; membrane-cytoskeleton linker *ANK2*^*51*^; microtubules-actin linker *MACF1*^*52,53*^, which also showed increased burden in astrocytes); and protein aggregate regulation (Aβ-downregulated *ATP5B*^*54*^; endosomal protein-aggregate mediator *ZFYVE16*; and glial-specific tauopathy inclusion mediator *CRYAB*^*55,56*^).

**Figure 4.**
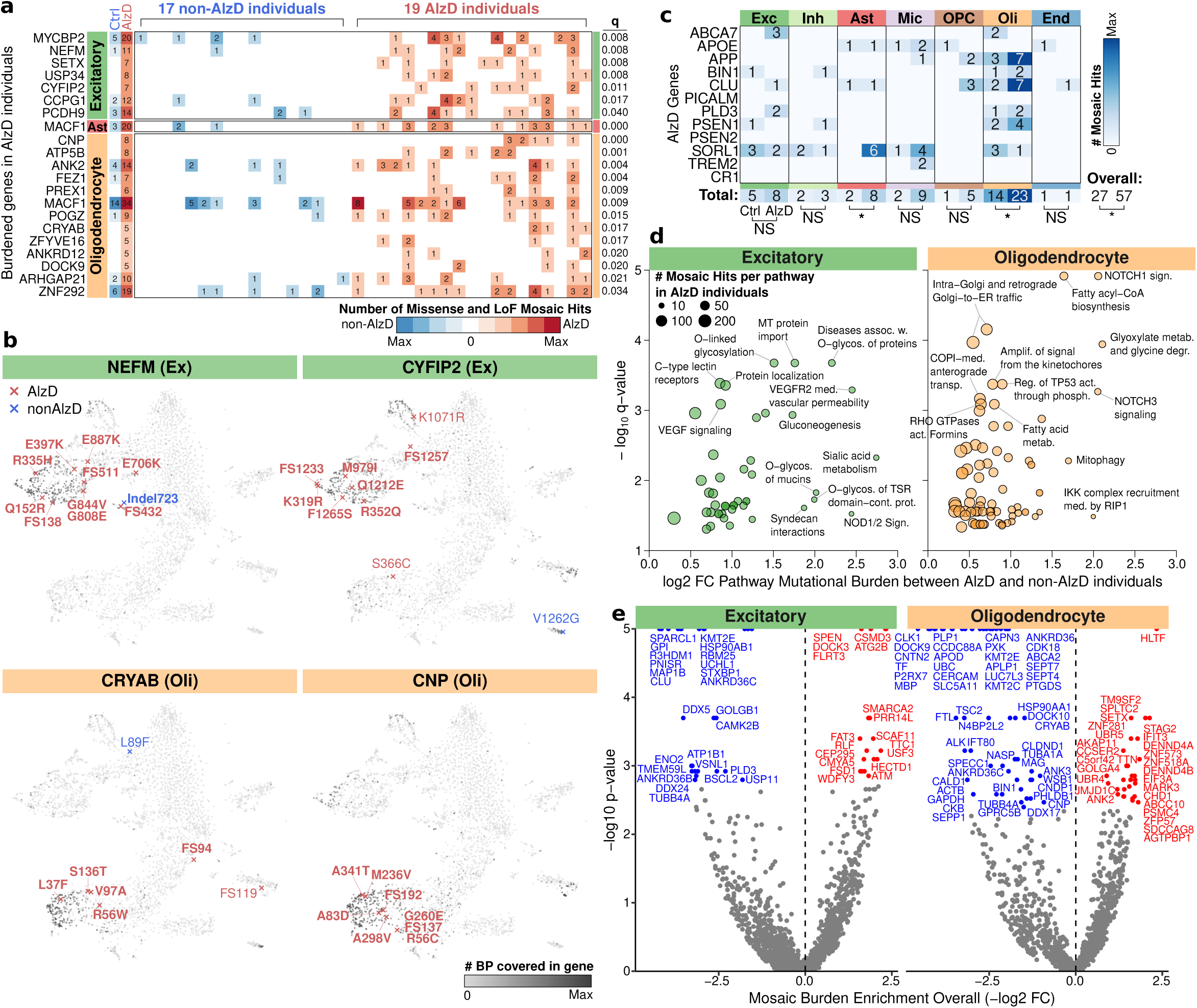
Gene-level and pathway-level mutational burden. **a**. Cell-type-specific mutational burden (counts) for AlzD (red) vs. non-AlzD (blue) individuals (columns) for increased-AlzD-burden genes (rows) (binomial q<0.05 FDR). **b**. Individual cells contributing to burden calculations in panel (a) for 4 example genes, with specific mutations labeled for AlzD (red) and non-AlzD (blue) individuals, for both AlzD-enriched-burden cells (bold) and all other cells (non-bold). Shade: gene-level nucleotide coverage. **c**. Mosaic hits (counts) in AlzD (right) vs. non-AlzD (left) individuals across cell types (column pairs) for known AlzD genes. Significance: Binomial q<0.05 FDR. **d**. Increased-AlzD-burden pathways (circles) in excitatory neurons (green) and oligodendrocytes (orange). X-axis: burden. Y-axis: -log_10_q-value. Size: mutation count in pathway. Labeled: q<0.001 or log2fc>1.5. **e**. Genes significantly enriched (red) or depleted (blue) for overall (AlzD-agnostic) mutational burden. X-axis: burden log-fold-change. Y-axis: -log_10_p-value. Labeled: q<0.05.

We next studied the individual cells where these mutations were found in the context of the tSNE gene expression landscape, pinpointing both the cell type and the cell state of each mutation-harboring cell for each gene, and the clinical status (AlzD vs. non-AlzD) of the individuals where the mutation was detected (**Fig. 4b, Supplementary Fig. S12-S14, Supplementary Table S2**). Indeed, we find that strong-effect missense and frame-shift mutations from these genes are concentrated in cells from relevant cell types, and that these cells come from AlzD individuals, consistent with the AlzD-specific and cell-type-specific enrichment of the mutational burden for these genes. Moreover, we find that the cells showing highest expression and highest coverage for these genes indeed match the cell types with AlzD-specific mutational enrichment, indicating that these enrichments are not an artifact of high-burden cells or senescent cells, which do not contribute any of the observed mutations. Even within excitatory neurons and oligodendrocytes, mutations in these genes are found in cells from sub-clusters corresponding to senescent-distal lower-burden cells, rather than the senescent-proximal higher-burden sub-clusters. These properties provide a strong affirmation of our discovery strategy, and indicate that these gene-enriched mutational enrichments are less likely to represent solely a consequence of dysregulation leading to increased mutation in these pathways.

We next searched for pathways that show greater mutational burden in AlzD individuals, even if individual genes within them did not rise to significance alone. First, we tested a set of 12 known AlzD genes and found significant mutational burden in AlzD, both overall (q=0.024) and in a cell-type-specific manner with burden concentrating in oligodendrocytes (23 mutations vs. 14, q=0.024) and astrocytes (8 vs. 2, q=0.025) (**Fig. 4c**), indicating that inherited-variant-containing genes are also recurrent targets of somatic mutations. Second, we tested 658 Reactome pathways with sufficient coverage for greater AlzD vs. non-AlzD mutational burden, and found 49 significantly-enriched pathways for excitatory-neuron mutations, and 80 significantly-enriched pathways for oligodendrocyte mutations (vs. only 19 and 4 pathways in controls) (**Fig. 4d, Supplementary Figure S15** and **Supplementary Table S3**).

For neuronal mutations, enriched pathways with greater AlzD burden included: neuron energy regulation (glycosylation, sialic acid metabolism); protein turnover and degradation (neddylation); DNA damage repair (excision repair); stress response (hypoxia response, syndecans, EPHA-mediated growth cone collapse); transport (mitochondrial import, protein localization, vesicle transport), and diverse signaling pathways. Oligodendrocyte mutations showed significantly greater AlzD enrichment in pathways involving: lipid metabolism (fatty acid metabolism); signaling (Notch1/3); endocytic vesicular transport (Golgi-to-ER retrograde, anterograde, endosomal sorting, clathrin-mediated, COPI); cell cycle (kinetochores); DNA damage (p53 phosphorylation); and proteostasis (HSP90 chaperones).

We also evaluated overall mutational burden in the aged human brain, across both AlzD and non-AlzD individuals, to search for genes with significantly-greater and significantly-lower burden. Greater-burden genes were enriched in: microtubule maintenance (*ATM, USP33, LYST, MARK1, CEP350, PCM1, BICD1, CTNNB1*) and synaptic cleft (*NLGN1, LGI1*) for neurons; DNA binding and gene expression regulation (*SETX, ZNF573, DENND4B, JMJD1C*) for glial mutations; and genomic stability, cytoskeleton dynamics and cell survival across cell types (**Fig. 4e, Supplementary Table S4, Supplementary Fig. S16-S17**).

Lower-burden genes were enriched in: axon targeting (*STXBP1, UCHL1*) and RNA binding (*COLGB1, HSP90AB1, PEBP1*) in neurons; cell-identity genes of myelination (*ABCA2, CNTN2, MBP, PLP1, MAG, MYO5A, CLU*) and microtubule-associated processes (*ATRX, CNTN2, HSPA1B, TUBA1A,GAPDH, TUBB4A*) in glia; and housekeeping and cell-identity marker genes across cell types. Mutational depletion might reflect a combination of decreased mutation rate across genomic regions, differential targeting of repair machinery proteins, or cellular-level selective pressure that deplete mutations leading to cell death.

Glia and neurons showed mutational enrichment and depletion in different genes, and their enriched-mutation pathway were largely distinct. However, they both showed mutational depletion in tau-regulating proteins (*CLU, GPI, HSP90AB1, STXBP1* in neurons, p_adj_=0.042; *CLU, HSP90AA1, ACTB, BIN1* in glia, p_adj_=0.019), with CLU, a GWAS^11^ and neuropathology^57^ AlzD-associated secreted chaperone that inhibits protein aggregation, showing mutational depletion in both.

## Discussion

Our study constitutes the first report of cell-type-specific mutational burden in affected-vs-unaffected individuals in AlzD, by leveraging single-cell full-transcript RNA sequencing technology (SMART-seq2) and matched whole-genome-sequencing to jointly infer somatic coding variants and their corresponding cell type and cell state, thus enabling us to understand the functional impact of mutational changes on genes and pathways.

This approach allowed us for the first time to partition somatic burden into the cell types and states where it occurs, and to survey the mosaic burden across multiple neuronal and glial cell types (while previous studies focused exclusively on neurons^27,58^). This revealed that overall mutational burden is higher in glial cells than in neurons, consistent with continued glial cell divisions in contrast to post-mitotic neurons. It also revealed that overall mutational burden, irrespective of disease status, concentrates in distinct genes and pathways between glial cells and neuronal cells, while other genes and pathways are depleted for mutations. Along with neuronal mutations that directly impact neuronal function, glial mutations can play important roles in brain changes, through indirect effects on neuronal functions via glial cells, and through direct effects on glial functions.

Our paper is also the first to report disease-enriched somatic mutational burden for a neurodegenerative disorder, where accumulation of somatic mutation burden across a lifetime of exposures is particularly relevant. We found that strong-effect somatic mutations in AlzD cluster in specific cell types, with excitatory neurons, astrocytes and oligodendrocytes emerging as primary drivers of AlzD-associated somatic mutations, highlighting the combined roles of neuronal cells, glial cells, and their interactions in AlzD^59,60^.

This greater mutational burden in AlzD also concentrated in specific genes and pathways, which differed between cell types. This included AlzD-associated genes and pathways from GWAS, including cholesterol metabolism, endosomal trafficking, and cytoskeletal systems. This also included AlzD-associated genes and pathways from rare variant and whole-genome-sequencing studies, including in protein aggregation, degradation, neurofilament tangles, and tau-associated proteins^61–63^. Our somatic burden analysis also captured novel pathways that were not previously genetically implicated, including oligodendrocyte myelination, post-translational modification, microtubule dynamics, and neuronal energy regulation through glycosylation, and stress response. These provide a new set of actionable targets for therapeutic development, which may have direct strong effects on neuronal function, but which previously lacked a genetic basis, as they lie outside the previously-studied range of the allele frequency spectrum from common to rare alleles.

In addition to elucidating the roles of neuronal and glial cell types in AlzD, our work highlights the importance of a new “senescent” cell population in the aging brain, which may represent a distinct cell type associated with aging, and was strongly enriched in AlzD. Even though these senescent cells are positioned in the two-dimensional tSNE embedding between neurons and oligodendrocytes, they show a distinct set of marker genes than either cell type, and instead might represent a collection of cells from different cell types that have lost their identity. As a group, senescent cells showed the largest mutational burden, both overall, and specifically in AlzD, which may have contributed to them losing their differentiated identity and entering a low-transcription and possibly pre-apoptotic state.

Our approach also allowed us to capture not only cell type identity, but also cell state, by relating the single-cell gene expression profile of each cell to the specific set of mutations it carries, and thus connect genotype to phenotype at single-cell resolution. Indeed, we saw a gradient of somatic mutational load along single-cell gene-expression space, indicating a strong correlation between the two, both between cell sub-clusters, and within each sub-cluster, indicating a continuum of mutational burden vs. expression dysregulation. For example, neuronal cells closest in gene expression to the senescent cells also showed increased mutational burden, while neuronal cells furthest in gene expression showed the lowest mutational burden. Similarly, oligodendrocyte gene expression sub-clusters correlated with mutational burden and distance from senescent cells. This concordance between accumulation of mutations and gene expression profiles at the cellular level may indicate that mutational burden accumulation results in phenotypic alterations; conversely, dysregulation at the single-cell level might also lead to increased mutations via DNA damage, decreased mismatch repair, and increased double-stranded breaks, which were previously-implicated in AlzD^64^.

It is challenging to distinguish whether mutational burden drives pathology, results from cellular dysfunction, or both. In the first case, high mutational burden can lead to cellular dysregulation, gene expression changes, and cellular pathology. Conversely, cellular dysfunction caused by other mechanisms (eg. protein aggregation, impaired protein turnover, DNA repair disruption) can lead to increased DNA damage and somatic mutation accumulation. Both processes may be acting simultaneously as well, with increased dysregulation leading to increased mutation accumulation and vice versa.

It is important to note that our approach only captures a fraction of the mosaic landscape that exists in aging and AlzD. First, we only annotate mosaicism in expressed transcripts, which has the advantage of focusing our sequencing efforts on strong-effect exonic mutations and on expressed genes, but does not capture non-coding regions, low-abundant transcripts, or alleles that lack expression. Second, RNA-based mutational inference can be biased due to transcription-associated mutations, transcription-associated repair, and the decreased stability of RNA relative to DNA, and thus is not well-suited for estimating an overall rate of somatic mutation; as such, we instead focus on relative differences here. Third, we do not capture the mosaic burden and cellular pathology incurred by cells that have already apoptosed, and even for senescent cells that exhibit low levels of transcription, our RNA-based approach likely misses many of their mutations in unexpressed segments of their genome.

Going forward, these limitations can be remedied in part by the use of technologies that simultaneously profile DNA and RNA from the same cell, and thus can decouple expression from genotype, and more directly ascertain how mosaic mutations relate with cellular state and gene expression levels. In addition, spatial transcriptomic approaches can enable direct comparison of cellular phenotypes in the context of surrounding pathology and their somatic mutations of neighboring cells.

Our study provides a foundational framework for analyzing such datasets, and initial insights on the relationship between somatic mutational burden, cellular state, and dysregulation of specific genes and pathways contributing to AlzD pathology. Our initial findings can be reinforced and refined as new technological advances make joint profiling of DNA and RNA from large numbers of cells commonplace in the study of the aging human brain in health and disease.

## Acknowledgements

We thank Alvin Shi, Jose Davila-Velderrain, Shahin Mohammadi, Khoi Nguyen, Michael Gutbrod, Jackie Yang, and all members of the Kellis and Tsai Labs at MIT and the Picower Institute for discussions and feedback. We thank the study participants and staff of the Rush Alzheimer’s Disease Center. This work was supported in part by NIH grants R01AG058002, RF1AG054012, U01NS110453, R01AG062335, UG3NS115064, RF1AG062377 (M.K. and L.H.T), R01AG067151, R01MH109978, U01MH119509, R01HG008155, U24HG009446 (M.K.), RF1AG054321 (L.H.T.), P30AG10161, R01AG15819, R01AG17917, U01AG46152, U01AG61356, RF1AG57473 (D.A.B.), the Cure Alzheimer’s Fund CIRCUITS consortium, and the Alana Foundation. H.M. was supported by an Early Postdoc Mobility fellowship from the Swiss National Science Foundation (P2BSP3_151885).

## Author contributions

M.Ko., C.B., and M.Ke. designed the study, carried out the analysis, and wrote the manuscript with help and input from all authors. Y.P.P. and S.S. provided help with computational and statistical analyses. H.M. and Z.P. carried out single-cell profiling. D.A.B. provided the human post-mortem brain samples. L.H.T. and M.Ke. supervised the study.

## Methods

### Single-nucleus SMART-seq2

#### Cohort and metadata

We assembled a cohort of 47 individuals, 22 with Alzheimer’s dementia and 25 age and sex matched persons with no-dementia from the ROSMAP cohorts^28,65^. The cases had a clinical diagnosis of Alzheimer’s dementia, mild cognitive impairment, or no cognitive impairment based on a detailed clinical evaluation as previously described^66,67^. For mosaic mutational burden analysis we additionally split the cohort by overall histopathology reported levels of β-amyloid load, PHFtau tangle density, and Braak stage, where stages 1-4 indicate the absence of NFTs in prefrontal cortex (PFC) and stages 5-6 indicate the presence of NFTs in the PFC (**Sup. Fig. 3A-G** and **Supplementary Table S5**)^68,69^.

#### Isolation of nuclei from frozen postmortem brain tissue for SMART-Seq2 plate-based single-nucleus RNA-sequencing

We adapted the protocol for the isolation of nuclei from frozen postmortem brain tissue from Swiech et al.^70^. All procedures were carried out on ice or at 4 °C. Briefly, postmortem brain tissue was homogenized in 2 ml Homogenization Buffer (320 mM Sucrose, 5 mM CaCl2, 3 mM Mg(Ac)2, 10 mM Tris-HCl pH 7.8, 0.1 mM EDTA pH 8.0, 0.1% IGEPAL CA-630, 1 mM b-mercaptoethanol, 0.4 U/microliter Recombinant RNase Inhibitor (Clontech)) using a Wheaton Dounce Tissue Grinder (10 strokes with the loose pestle). 3 ml of Homogenization Buffer was added (final volume 5 ml) and the homogenized tissue was incubated on ice for 5 minutes. Then the homogenized tissue was filtered through a through 40 μm cell strainer, mixed with an equal volume of Working Solution (83% OptiPrep™ Density Gradient Medium (Sigma-Aldrich), 5 mM CaCl2, 3 mM Mg(Ac)2, 10 mM Tris HCl pH 7.8, 0.1 mM EDTA pH 8.0, 1 mM β-mercaptoethanol) and loaded on top of an OptiPrep density gradient (10 ml 29% OptiPrep solution (29% OptiPrep™ Density Gradient Medium, 134 mM Sucrose, 5 mM CaCl2, 3 mM Mg(Ac)2, 10 mM Tris-HCl pH 7.8, 0.1 mM EDTA pH 8.0, 1 mM β-mercaptoethanol, 0.04 % IGEPAL CA-630, 0.17 U/microliter Recombinant RNase Inhibitor) on top of 5 ml 35 % OptiPrep solution (35% OptiPrep™ Density Gradient Medium, 96 mM Sucrose, 5 mM CaCl2, 3 mM Mg(Ac)2, 10 mM Tris-HCl pH 7.8, 0.1 mM EDTA pH 8.0, 1 mM β-mercaptoethanol, 0.03 % IGEPAL CA-630, 0.12 U/microliter Recombinant RNase Inhibitor)). The nuclei were separated by ultracentrifugation using an SW32 rotor (20 minutes, 9000 rpm, 4 °C). 3 ml of nuclei were collected from the 29%/35% interphase and washed with 15 ml ice cold PBS containing 0.5% BSA and 2 mM EDTA. The nuclei were centrifuged at 100 g for 5 minutes (4 °C) and resuspended in 1 ml PBS containing 0.5% BSA and 2 mM EDTA. Then the nuclei were stained by adding two drops of NucBlue Live ReadyProbes Reagent (ThermoFisher Scientific, catalog number: R37605) and passed through a 40 μm cell strainer (Falcon Cell Strainers, Sterile, Corning, product 352340). Single Hoechst-positive nuclei were sorted into 96-well plates (Eppendorf twin.tec PCR Plate 96, catalogue number 951020401) containing 5 microliters of Buffer TCL (QIAGEN, catalogue number 1031576) per well containing 1% beta-mercaptoethanol (Sigma-Aldrich, catalogue number M6250), snap frozen on dry ice, and then stored at −80 °C before whole transcriptome amplification, library preparation and sequencing.

#### Reverse transcription, whole-transcriptome amplification, library construction, sequencing

Single-nucleus RNA sequencing libraries were generated based on the SMART-Seq2 protocol^71^ with the following modifications. RNA from single cells was first purified with Agencourt RNAClean XP beads (Beckman Coulter, catalogue number A63987) before oligo-dT primed reverse transcription with Maxima H Minus Reverse Transcriptase (Thermo Fisher, catalogue number EP0753), which was followed by 21 cycle PCR amplification using KAPA HiFi HotStart ReadyMix (KAPA Biosystems, catalogue number KK2602) with subsequent Agencourt AMPure XP bead (Beckman Coulter, catalogue number A63881) purification. Libraries were tagmented using the Nextera XT DNA Library Preparation Kit (Illumina, catalogue number FC-131-1096) and the Nextera XT Index Kit v2 Sets A,B,C, and D according to the manufacturer’s instructions with minor modifications. Specifically, reactions were run at one fourth the recommended volume, the tagmentation step was extended to 10 minutes, and the extension time during the PCR step was increased from 30 seconds to 60 seconds. Libraries from 192 cells with unique barcodes were combined and sequenced on the Illumina HiSeq 2000 platform at the MIT BioMicro Center.

#### Read processing

We aligned, uniformly-processed, and called mutations on each of 6,180 single nuclei from 47 individuals (average of 131.5 per individual) following GATK best practices pipeline for RNA-seq^72^. We first aligned 40-bp single-end reads from each sequenced single nucleus separately with the STAR aligner^73^ against the b37 genome with decoy contigs using a two pass alignment (options: --outFilterMultimapNmax 20 - -alignSJoverhangMin 8 --alignSJDBoverhangMin 1 --alignIntronMin 20 --alignIntronMax 1000000). We then used Picard tools^74^ to revert and merge the alignment with unaligned reads and marked duplicates on the merged bam file. We identified and removed alignments on decoy contigs, sorted and fixed NM, MD, and UQ tags with Picard tools, filtered duplicates, unmapped, and non-primary alignment reads, and split reads by Ns in their CIGAR string. Finally, to generate a bam file for transcriptomics and mutation calling we ran two passes of GATK^72^ BaseRecalibrator for each cell against the b37 genome with decoy contigs, using dbSNP^75^ v138 and the Mills 1000G indels^76^.

### Transcriptional profiles

#### Cell identities

For each of 6,180 cells, we used HTSeq-count^77^ on its filtered and recalibrated bam file to compute the cell’s transcriptomic coverage over each gene’s exons in GENCODE gene annotation (v28 lifted to b37). We used SCANPY^78^ to process and cluster the expression profiles and infer cell identities. We kept only 19,765 protein coding genes detected in at least 3 cells and filtered out 37 cells with less than 100 expressed genes. We identified and filtered out 322 additional cells showing very strong individual-specific batch effects, leaving 5,821 cells over 47 individuals. We used the filtered dataset to calculate the low dimensional embedding of the cells (t-Stochastic Neighbor Embedding: t-SNE) (default parameters, perplexity=50), built a nearest neighbors graph (n=10), and clustered it with the Louvain clustering (resolution=2), giving 24 preliminary clusters. We then manually assigned clusters based on the following 2-3 major marker genes per class: Neuronal: *GRIN1, SNAP25, SYT1*; Excitatory neurons: *CAMK2A, NRGN, SLC17A7*; Inhibitory neurons: *GAD1, GAD2*; Astrocytes: *AQP4, GFAP*; Microglia: *C3, CD74, CSF1R*; Oligodendrocytes: *MBP, MOBP, PLP1*; Oligodendrocyte progenitor cells (OPCs): *PDGFRA, VCAN*; Endothelial: *FLT1* and *CLDN5*. We merged clusters sharing marker genes to obtain 9 final clusters, defining two broad neuronal subtypes (1,170 excitatory and 221 inhibitory cells), four glial clusters (220 astrocytes, 255 microglia, 2121 oligodendrocytes, and 94 OPCs), 40 endothelial cells, 400 cells with strong individual-specific batch effect and cancer signatures (*DNMT3A, COL6A3*), and 1,300 senescent cells, marked by *CARD8, FAM126A, IRX2, ALK, SENP7*, and *GMFB* and lower overall transcription (**Supplementary Figure S8 and Supplementary Table S6**). In burden analyses, where we used the 36 individual cohort, we kept 4,440 cells corresponding to these individuals (Ex: 935, Inh: 167, Ast: 159, Mic: 188, OPC: 74, Oli: 1,382, End: 32, Batch: 400, Sen: 1,103). As a final step of QC, we removed the 400 cells showing batch effect. In our previous study^28^, we did not observe a senescent cell population, likely due to the protocol’s selection bias for barcoded cells with higher numbers of UMIs.

#### Sub-clustering

We further sub-clustered the cell types with the largest number of cells (N ≥ 1,000, excitatory neurons, oligodendrocytes, and senescent clusters) by Louvain clustering (resolution=0.3). To further define the clusters and sub-clusters, we performed marker-gene discovery on the clusters against all cells and on the sub-clusters against their overall cluster with scanpy, removing unannotated and MT-like genes. We show the top 10 marker genes per cluster in **Fig. 2F** and per subcluster in **Sup Fig. 10A-C**.

#### Covariate enrichment

We calculated the enrichment of each level of each of the major covariates (sex, cognitive impairment level, cognitive impairment score, NFT level, amyloid plaque level, braak stage, and presence/absence of NFTs in the pre-frontal cortex) in each cluster and sub-cluster by the hypergeometric test, and show enrichments with -log_10_p > 2 (**Fig. 2E and 3F**).

### Mutation calling

#### WGS

For 37 of the 47 individuals, we obtained germline variants from ROSMAP (at https://www.synapse.org/#!Synapse:syn10901595)^31,65^. Variants were profiled using whole genome sequencing (WGS) at 30x coverage with 150bp paired-end reads, aligned on the GRCh37 genome, and processed using the GATK best practices pipeline, including marking duplicates, indel realignment, and base quality recalibration, and called using the HaplotypeCaller and GenotypeGVCFs^72^. To create a germline list to filter against our calls, we conservatively retained all variants with a depth of at least 3 and a non-reference allele fraction of at least 20% using vcftools. Of the 37 WGS datasets, 17 were performed on the prefrontal cortex, 11 on another brain region, and 9 on blood.

#### snRNA-seq

In order to call potential mosaic hits, we ran GATK’s HaplotypeCaller on each cell independently, merged all vcfs using GenomicsDBImport on each chromosome, ran GenotypeGVCFs on the merged cells in each individual using dbSNP v138 as a database of known SNPs, and concatenated vcfs with GatherVcfs^72^. Using GATK’s VariantFiltration, we filtered out any variants corresponding to the individual’s germline calls or in low-complexity regions^79^. We compared known germline variants to potential mosaic variants in the sn-RNA-seq data to establish additional variant filters and kept variants with QD > 15 at DP = 3, QD > 18 at DP= 4, QD > 20 at DP=4, and with FS < 10, SOR > .5, MQ < 61, or DP < 3 (**Sup. Fig 1A-C**). We kept any variants with over 20% allelic balance that also overlapped exons of protein coding genes in the GENCODE annotation, resulting in 66,664 variants across all cells (keeping 24.3% of 273,508 preliminary calls). We further filtered variants by flagging: **(a)** sites in blacklisted regions (chrY, chrM, decoy contigs, and scaffolds), or **(b)** in blacklisted genes (HLA locus, IgG genes, and pseudogenes); **(c)** RNA editing sites from RADAR v2, DARNED, or REDIportal^80–82^; **(d)** sites that appeared in more than 1 individual and potentially represented alignment artifacts; **(e)** sites in cells filtered out in the final transcriptome analysis; **(f)** calls with only one alternate supporting read; **(g)** 326 calls matching variants in ExAC with MAF > 1%^83^ (**Sup. Fig. 1D)**; **(h)** sites from 31 hyper-mutated flagged cells; and **(i)** merging calls found within 1-3bp of each other in the same samples into single indels (reduced 3,105 to 1,459 calls). This resulted in 56,608 (84.9% from 66,664 and 20.7% of 273k) final, filtered variants in 37 individuals with matched WGS, with mutations in 3,044 of 5,821 kept cells) for downstream analysis (**Sup. Fig. 1A-E**). For all mosaic burden analyses, we kept 55,447 variants (97.9%) from the 36 individuals with WGS and cognitive impairment scores (cogdx variable) between 1 and 5.

#### Variant annotation

We annotated variants with SnpEff v4.3^84^ using the GENCODE v28lif37 b37 genome annotation^85^. We grouped variants into synonymous (12,166, 21.5% of 56,608 variants from the 37 individuals), missense (29,071, 51.3%), loss of function (LoF, 13,623, 24.1%), and other variants (1,748, 3.1%). LoF variants included frameshift, start or stop variants, indels, and splice variants (**Sup. Fig. 2A-C**). We used ExAC to annotate the MAF of 3,636 rare matching variants (6.42% of filtered calls), with 3,283 (90.3%) at less than 0.01% and 1,880 (51.7%) at less than 0.001%^83^ (**Sup. Fig. 1D**). Finally, we annotated 53,789 (95%) variants with CADD scores^86^, of which 5,861 (10.9%) have scores up to 10, 13,484 (25.1%) between 10 and 20, 24,007 (44.6%) between 20 and 30, and 10,437 (19.4%) above 30 (**Sup. Fig. 2D**).

#### Mutational signatures

We computed the observed trinucleotide signatures and normalized signatures within each sample by enumerating the possible DNA tri-nucleotide contexts in genomic locations covered by snRNA-seq at a callable depth for each cell. We estimate and report the average % for each type of single-nucleotide variant (SNV) for all 36 individuals as **Figure 1E** and split by case/control status according to cognitive diagnosis in **Sup. Fig. 5A, 5B** (clipped at 10%). Mutational signature decomposition was performed relative to the COSMIC^38^ mutational signatures using deconstructSigs^87^ in R (**Sup. Fig. 5C**).

### Mutational Burden

#### Individual-level burden

We performed all mosaic mutational burden analyses on the 36 of the 37 individuals with WGS and mosaic variant calls. For burden analyses, we removed one individual of the 37 who had a cognitive score of 6 (other dementia). The 36 individuals contained 19 cases and 17 controls by cognitive diagnosis (scores of 1-3 vs. 4-5). Amyloid level split the burden cohort 20 to 16 (amyloid > 1), tangles split 15 to 21 (tangles > 5), and the presence of tangles in PFC split 14 to 22 (braaksc 1-4 vs. 5-6). For each individual, we estimated the average mosaic mutational burden as the number of mosaic calls divided by the number bases covered at a callable depth in the snRNA-seq data. We compared the distribution mosaic burden in cases to controls using a one-sided t-test. For variant type and CADD divisions, we computed burden estimates per individual per variant class and compared cases and controls, as defined by cognitive diagnosis, within synonymous, missense, and LoF variants, as well as in each CADD score bin.

#### Cell-level burden

For the cell-level analysis, we estimated the mutational burden for each individual in each cell type and in all neurons (label “Neuronal”: excitatory and inhibitory neurons) and all glia (label “Glial”: aggregating astrocytes, microglia, OPCs, and oligodendrocytes). We used one-sided t-tests to determine significance both between major cell groups (neuronal to glial and to senescent and glial to senescent), and within cell-types between cases and controls as defined by cognitive diagnosis. Additionally, we evaluated the effect of sex (male/female) on mosaic mutational burden both over all cell types in aggregate and at the cell-level by two-sided t-tests (**Sup. Fig. 8**). To evaluate the effect of age-of-death in a cell-type-dependent manner, we fit a linear regression to the per-individual burden averages within either neuronal (p=0.608) or glial cells (p=0.127). We also evaluated the interaction of age and cell-type by comparing the additive model to a model with the interaction term (comparison by anova, p=0.55).

### Gene and pathway enrichment

#### Gene-level mosaic burden

First, to discover genes significantly over-burdened in each cell type, we analyzed each major cell type separately, and tested each gene by comparing its mutational burden to that of matched random samples from a stratified full dataset. To stratify the dataset, we calculated the number of covered bases in the exons of each gene in each individual cell, annotated each covered gene in each cell with its coverage and number of hits as one case. We kept only protein-coding mutations annotated by SnpEff as missense or LoF variants. We reduced our dataset to 41,755 missense and LoF variants from 4,014 cells from the 36 individuals with paired WGS that were not filtered out in mosaic calling or transcriptome QC. Within each major cell type (and jointly for all neurons, all glial cells, and all cells), we created a stratified dataset by stratifying by sex (male/female), age (17 individuals up to 86.9 and 19 greater than 86.9), and by binning each cell by gene’s (sample’s) coverage. We tested all genes with at least one mutation and 10,000 bp of coverage in cases and controls. For each tested gene, we sampled 10,000 null genes from the stratified dataset, matching the gene’s coverage in each cell, and compared the gene’s burden to that of null examples to obtain a significance p-value. We used the qvalue^88^ R package to perform FDR correction. Per-cell-type pathway enrichment was performed using gprofiler^89^ for an over-representation test for genes with q-value < 0.05 in each cell type (**Supplementary Table S3**).

#### Gene-level mosaic burden in disease by cell type

Second, to discover genes that were significantly over-burdened in disease, we compared the relative rates of mosaic events in AD individuals versus controls as defined by clinical ascertainment of cognitive impairment. Within each cell type (and jointly for all neurons, all glial cells, and all cells), we tested all genes with at least 10,000 bp of coverage both in AD cases and controls and with at least one mutation in cases. We performed a two-sided exact binomial test for the number of mutations in the number of covered bp in cases against the estimated rate of mutation in controls, adding a pseudocount of one mutation to both cases and controls. FDR correction was performed using the qvalue^88^ R package. We also performed a permutation test as a secondary test for over-burdened genes in AD individuals. We performed the stratification as in the case of the disease-agnostic analysis, further stratifying the dataset by the estimated burden in each gene, agnostic to disease status (binned into 0-10^−5^, 10^−5^-10^−4^, 10^−4^-10^−3^, and 10^−3^ or greater). We tested all genes flagged as significantly over- or under-burdened by the binomial test. For each tested gene, we sampled 10,000 matched null examples from the stratified dataset and compared the overall ratio of hits in AD individuals to controls to the ratios from null examples in order to obtain a significance p-value (**Supplementary Table S4**). We evaluated the enrichment of mutations for bona-fide AD genes (*ABCA7, APOE, APP, BIN1, CLU, CR1, PICALM, PLD3, PSEN1, PSEN2, SORL1*, and *TREM2*) in aggregate in each cell type and overall using a binomial test and FDR correction as above.

#### Pathway-level burden in disease by cell type

We evaluated pathway-level differences in mosaic mutational burden in cases versus controls on the Reactome^90^ pathway maps obtained from the reactome.db R package^91^. In each cell type and in the aggregated cell types, we aggregated mosaic mutations and coverage from the gene-level to the pathway-level and kept all pathways with least 10,000 bp of coverage both in AD cases and controls, at least one mutation, between 25 and 250 genes, and where at least 90% of the pathway genes are expressed in the tested cell type. To compare each pathway’s rate of cases to controls, we performed a binomial test followed by FDR calculation for multiple testing correction as in the gene-level analysis (**Supplementary Table S5**).

## Supplementary Tables

**Table S1**. Gene-level AlzD-specific enrichments

**Table S2**. Burdened Genes Hits

**Table S3**. Pathway-level AlzD-specific enrichments

**Table S4**. Overall enriched and depleted genes

**Table S5**. Individual-Level Metadata

**Table S6**. Cell-Level Metadata

