## Supplementary Figures for "Single-cell mosaicism analysis reveals cell-type-specific somatic mutational burden in Alzheimer’s Dementia"

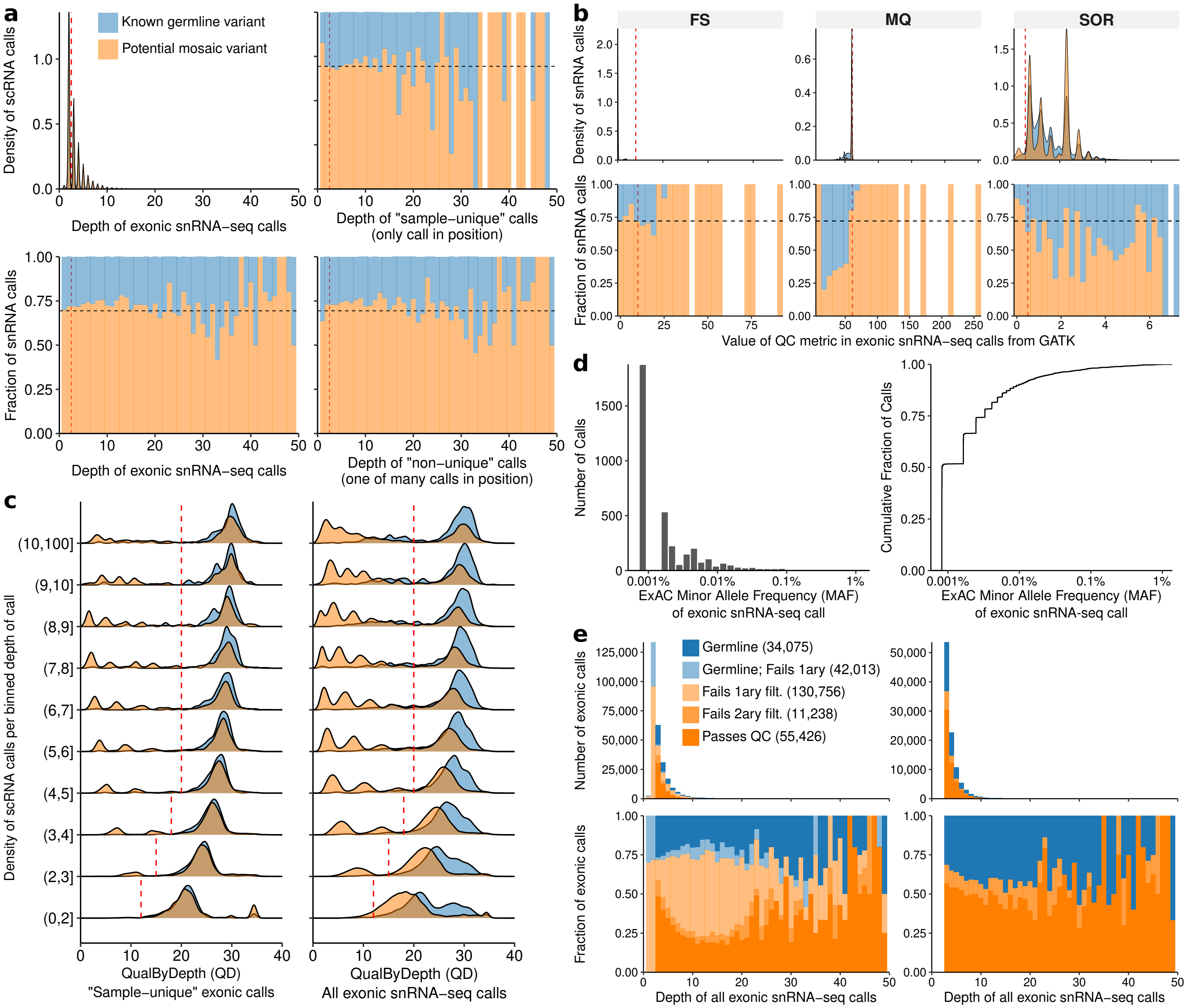


**Supplementary Figure 1. Variant filtering. a.** Distribution of calls from snRNA-seq by depth of call (DP), split into known germline variants by intersection with WGS or potential mosaic variants. Top-left: density of calls by depth; bottom-left: stacked histogram showing relative fraction of germline and potential mosaic calls per depth; top-right and bottom-right: breakdown of calls into sample-unique calls, where the call is supported across multiple samples (cells) in the individual versus non-unique calls. Horizontal line labels average fraction of potential mosaic calls relative to all calls. **b.** Comparison of FS (FisherStrand), MQ (RMSMappingQuality), and SOR (StrandOddsRatio) variant filtration metrics between known germline calls and potential mosaic calls in order to establish filtration cutoffs (vertical lines). **c.** Comparison of quality by depth (QD) metric at 10 depth cutoffs between validated calls corresponding to germline versus potential mosaic calls. As QD is evaluated on all samples in one individual and many germline variants are found across multiple cells, cutoffs are established on sample-unique calls, for which the germline distribution is comparable to that of potential mosaic events. **d.** Minor Allele Frequency (MAF) distribution of calls found in ExAC with MAF <1% (left) histogram of number of calls by MAF (right) cumulative distribution of all calls by MAF, where over 50% are at less than 0.001% MAF. **e.** Breakdown of calls by overall number (top) and relative fraction (bottom) including all calls (left) or just those passing the 1st round of filtering (right). 55,426 variants pass QC (in addition to 1,459 indel variants from merging calls, not shown).

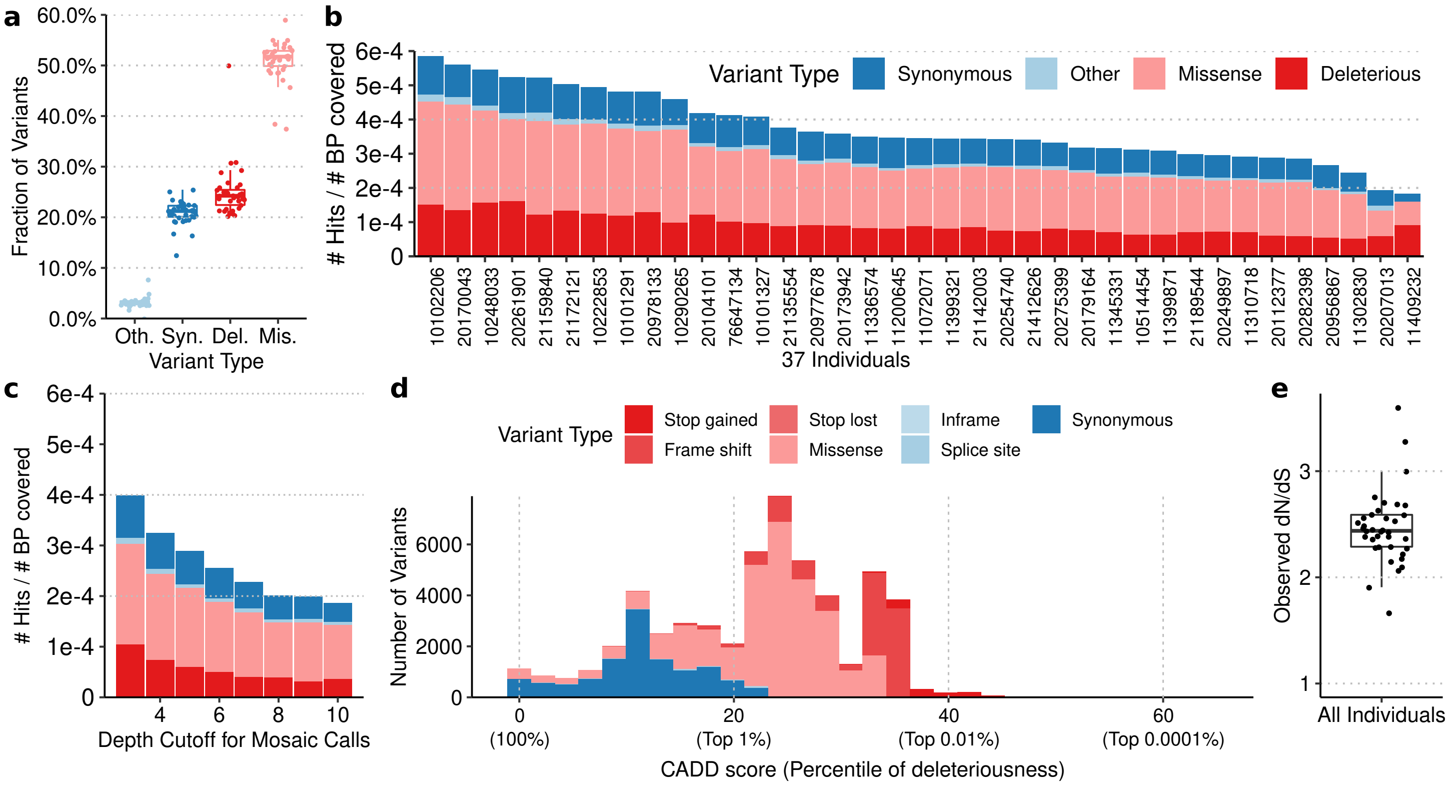


**Supplementary Figure 2. Mutational burden by variant type. a.** Distribution of variant type breakdown at the individual level, where each point is one individual, for missense (average: 51.3%), synonymous (21.5%), LoF (24.1%), and other (3.1%) mosaic variants. **b.** Overall mutational rate (number of mosaic hits over the number of bases covered at a callable depth) by individual and variant type. **c.** Overall mutational rate by depth of mosaic call and variant type. **d.** Distribution of CADD score against variant type for mosaic calls. **e.** Distribution of dN/dS rates across individuals.

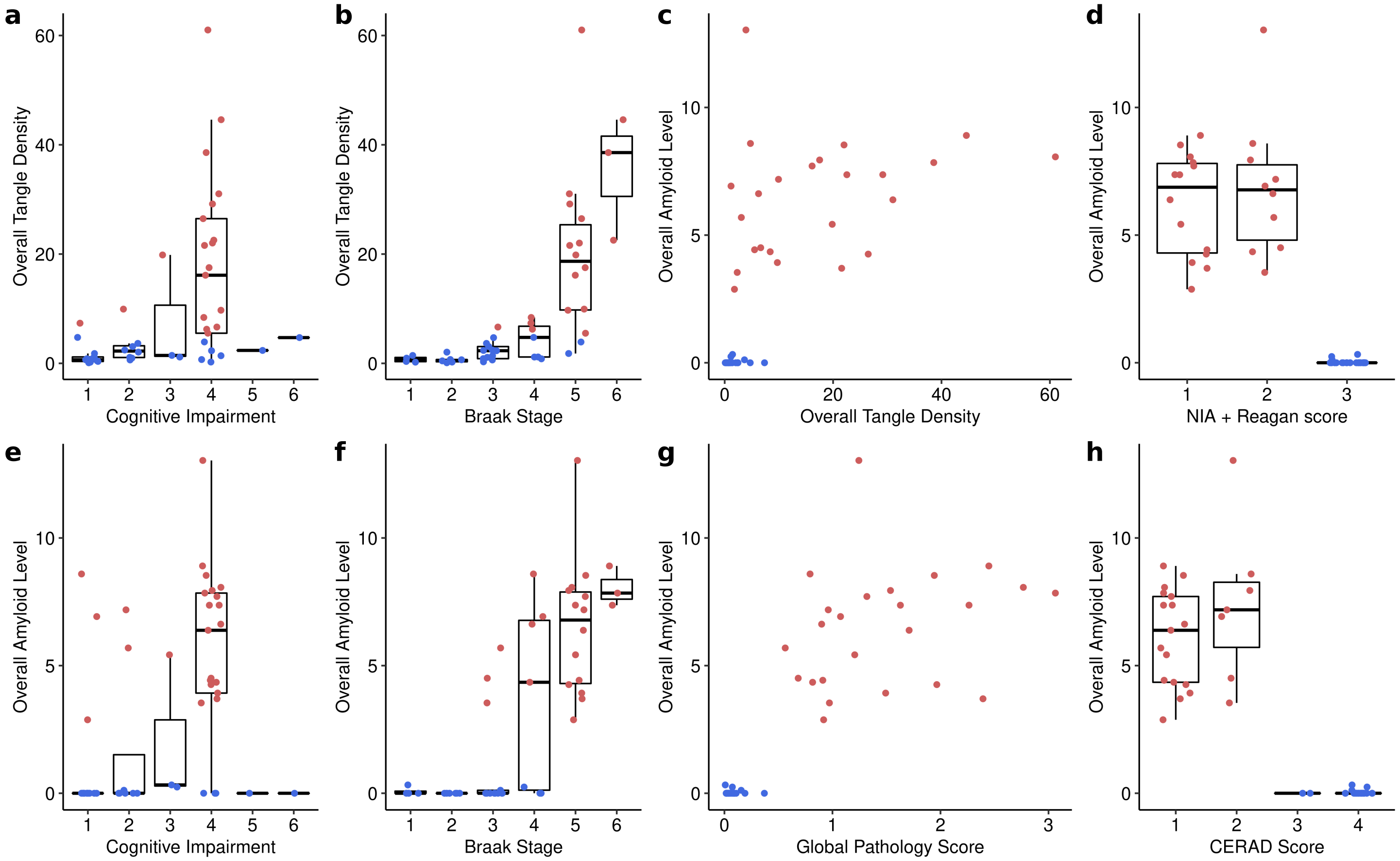


**Supplementary Figure 3. Cohort Ascertainment Measures.** Comparison of Alzheimer's dementia and disease ascertainment measures over the full 47 individual cohort used for transcriptional analysis. **a-b.** Comparisons of cognitive impairment (CI) stage and Braak stage against overall tangles levels (colored by high tangles level in red, low in blue) c-h. Comparisons against overall amyloid level, colored by amyloid level (high in red, low in blue) of overall tangle density, NIA/Reagan score, CI stage, Braak Stage, global pathology score, and CERAD score.

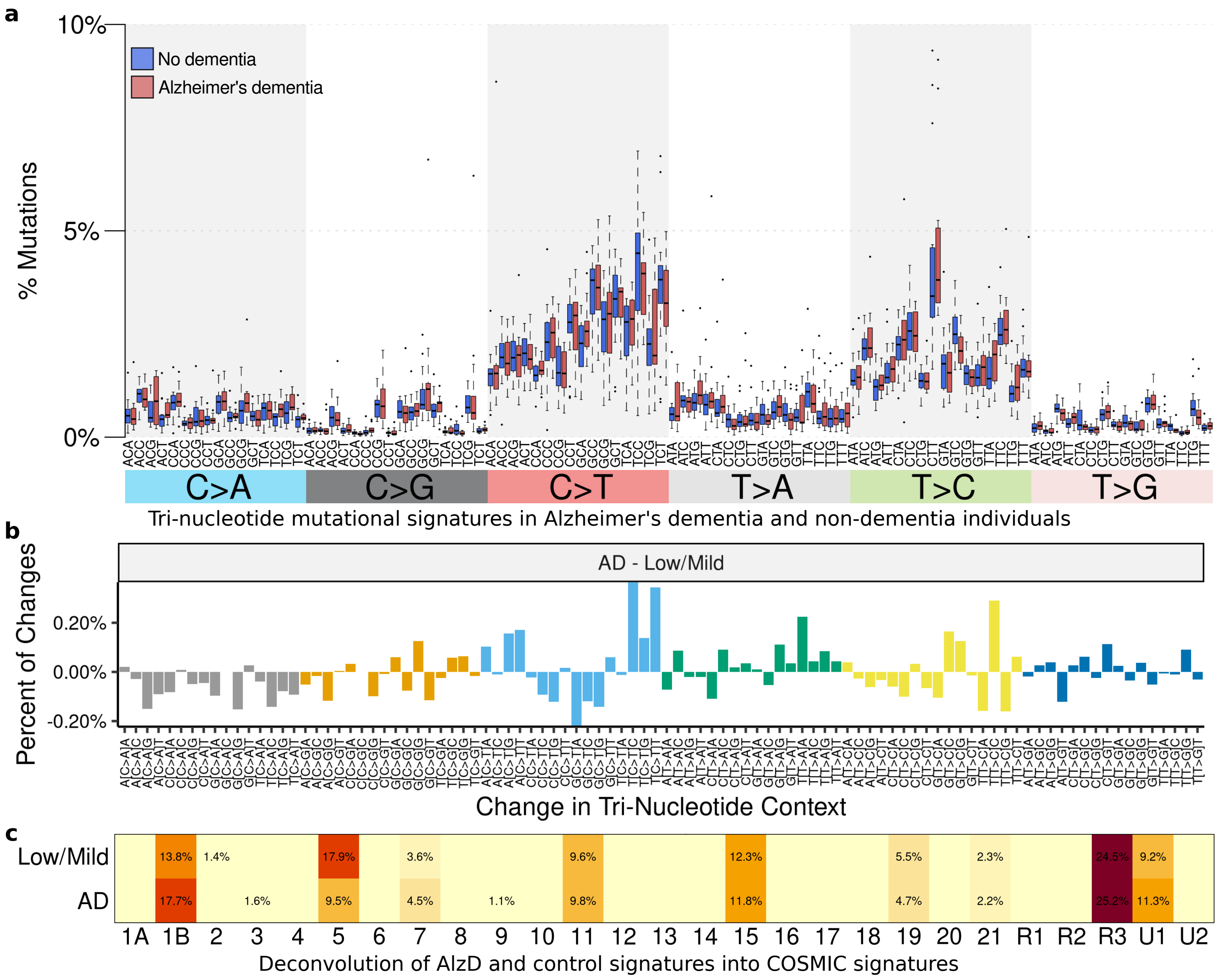


**Supplementary Figure 4. Mutational signatures in disease. a.** Distribution of mutational signatures by cognitive impairment status (non-dementia in blue and AlzD in red), where each point is an individual. No significant differences were found for any signature after p-value adjustment for multiple testing. **b.** Difference in the overall mutational signatures between Alzheimer's dementia and non-dementia individuals. **c.** Mutational signature deconvolution of the overall dementia and non-dementia signatures (rows) according to the COSMIC v1 signatures (columns).

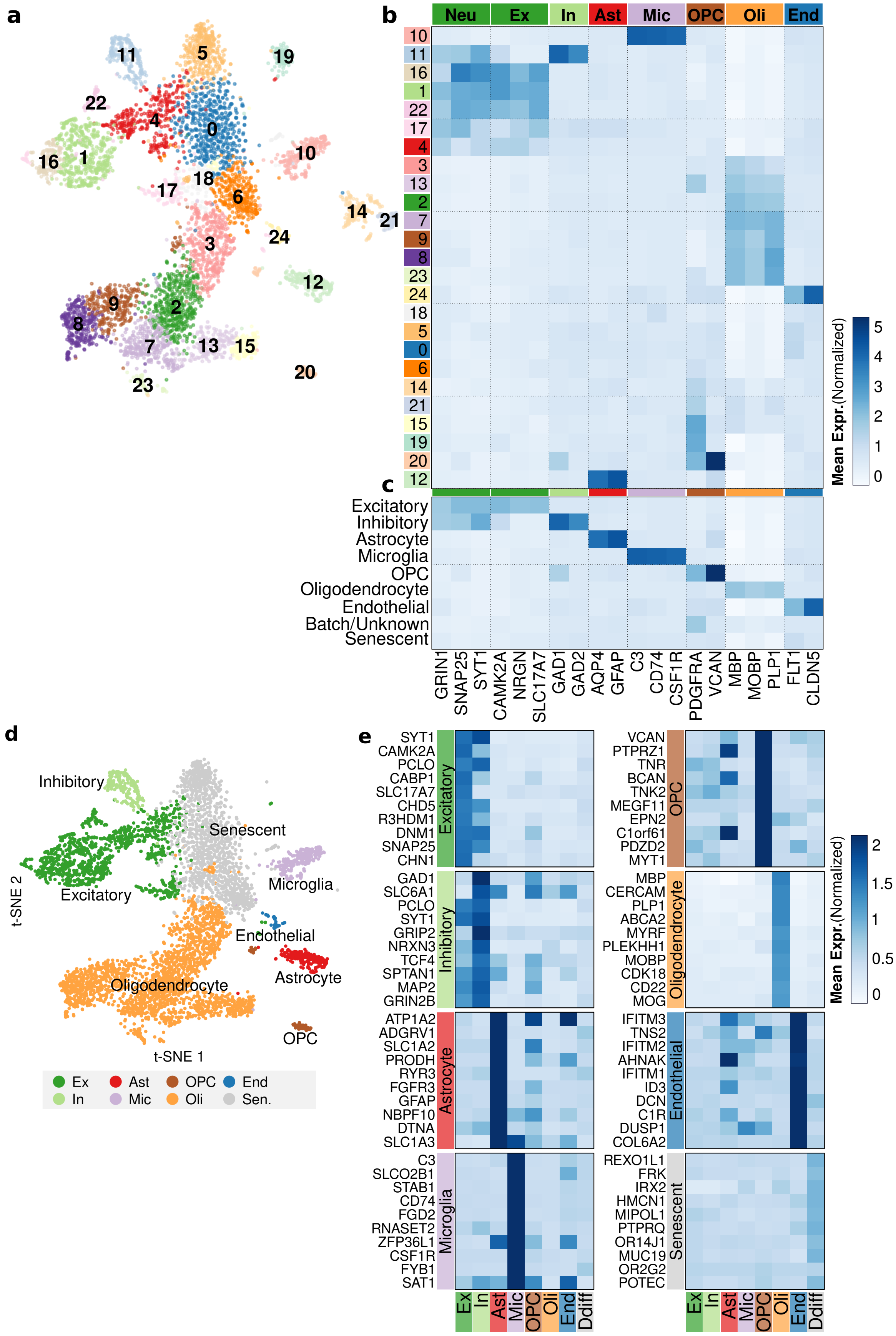


**Supplementary Figure 5. Cell type identification. a.** Preliminary louvain clusters for the t-SNE of 5,821 cells. **b.** Comparison of known brain cell type markers to louvain clusters. Sub-clusters 14 and 19-21 are removed in further analysis. **c.** Comparison of marker genes to cell types merged from louvain clusters. **d.** t-SNE plot for 5,421 cells from 47 individuals across 8 merged cell clusters (1,170 excitatory and 221 inhibitory cells), four glial clusters (220 astrocytes, 255 microglia, 2121 oligodendrocytes, and 94 OPCs), 40 endothelial cells, 400 cells with strong individual-specific batch effect and cancer signatures (DNMT3A, COL6A3), and 1,300 senescent cells. **e.** Learned marker genes for each cell type. The top 10 markers are shown for each cell type against their average expression from each cluster.

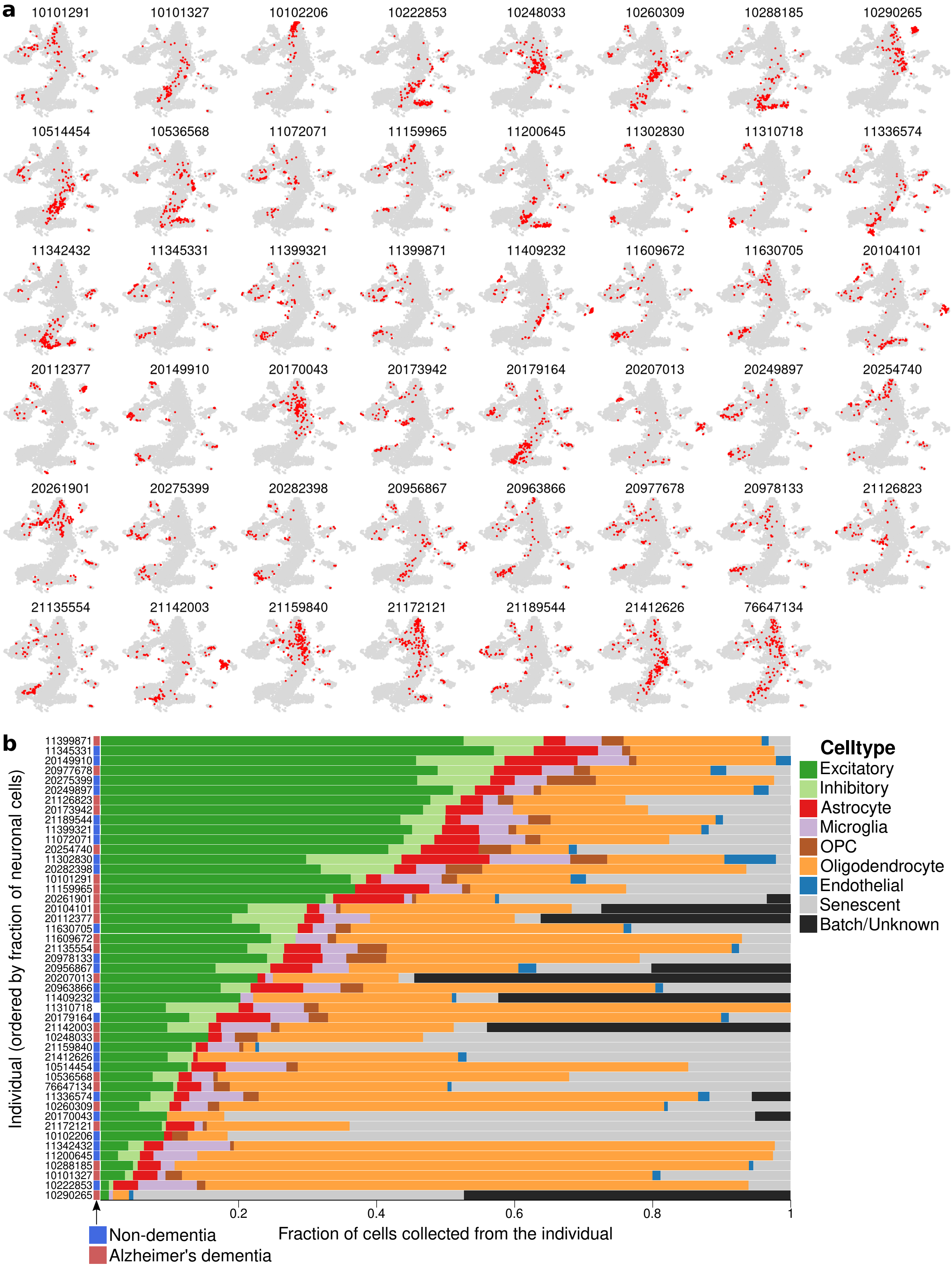


**Supplementary Figure 6. Cell type breakdown by individual. a.** Locations of each of the 47 individuals' cells on the t-SNE of 5,821 cells before removal of batch cells. **b.** Cell fractions per individual for 8 major cell types and one batch. Individuals are labeled as Alzheimer's dementia or non-dementia as ascertained by their cognitive impairment status.

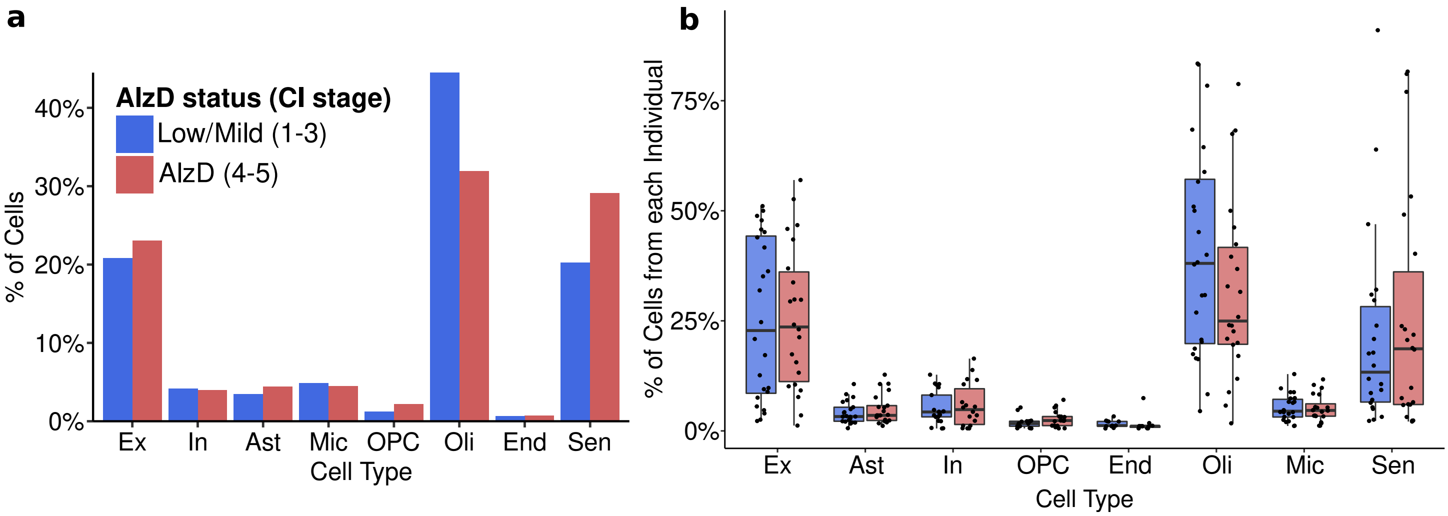


**Supplementary Figure 7. Cell fractions by disease status. a.** Overall percentage of cells from all AD individuals (22, cogdx in 4-5, red) and control individuals (24, cogdx in 1-3, blue) coming from each cell type. One of the 47 individuals has cogdx level 6 and is not included. **b.** Per-individual fraction of each cell type.

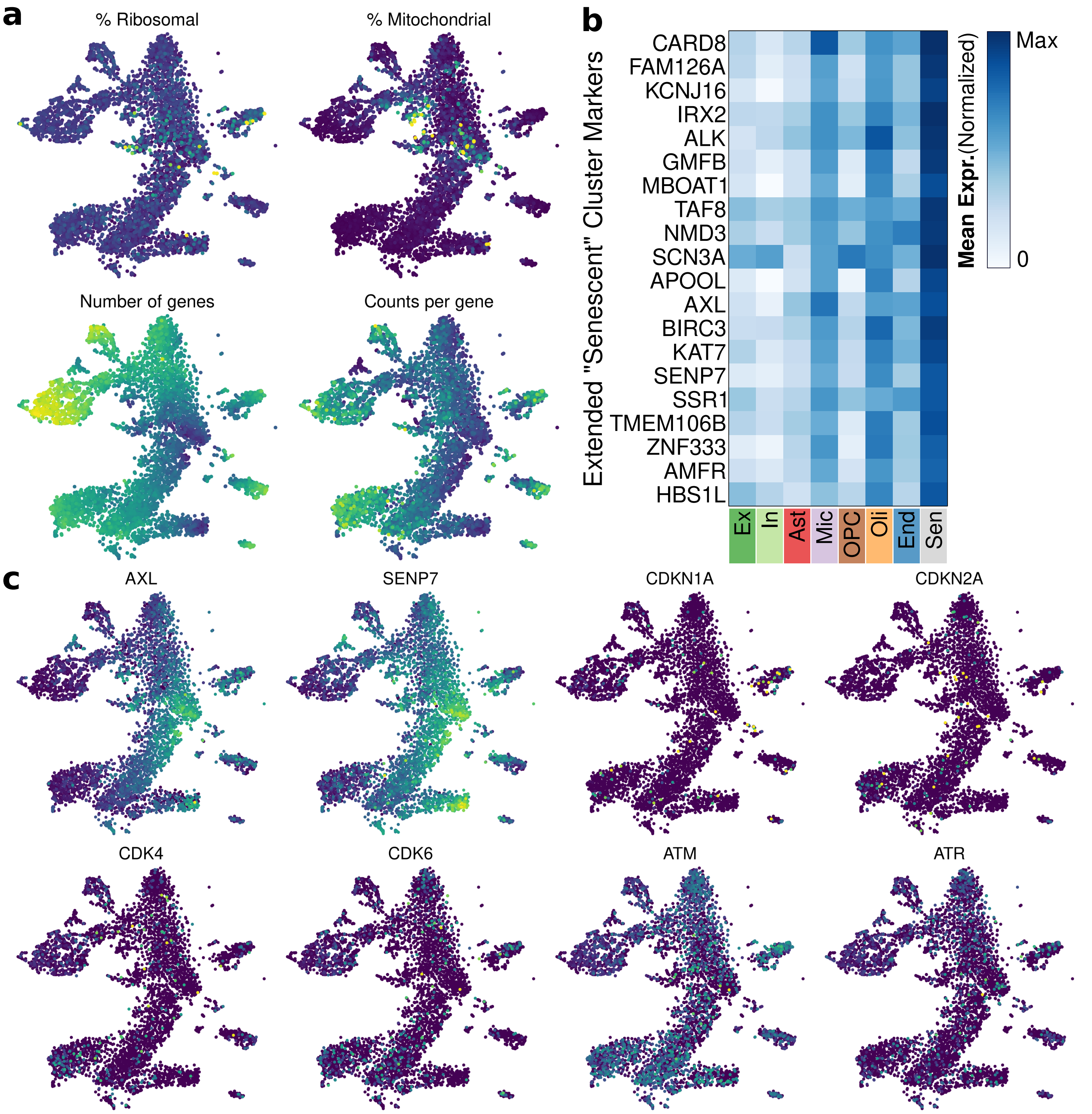


**Supplementary Figure 8.** Extended markers for t-SNE and "senescent" cluster. **a.** Percentage of reads from ribosomal or mitochondrial genes, number of genes, and number of reads per gene in each cell. "Senescent" cluster cells show overall lower expression and less counts per gene. **b.** Top 20 "senescent" cluster markers where at least half of the cells show greater than average expression of the gene. Marker genes were computed for the cluster against all cells by a wilcoxon test. **c.** Classical senescence markers, including CDKN1A and CDKN2A, which regulate CDK4 and CDK6, show low signal across the cells in the t-SNE (dark = low expression).

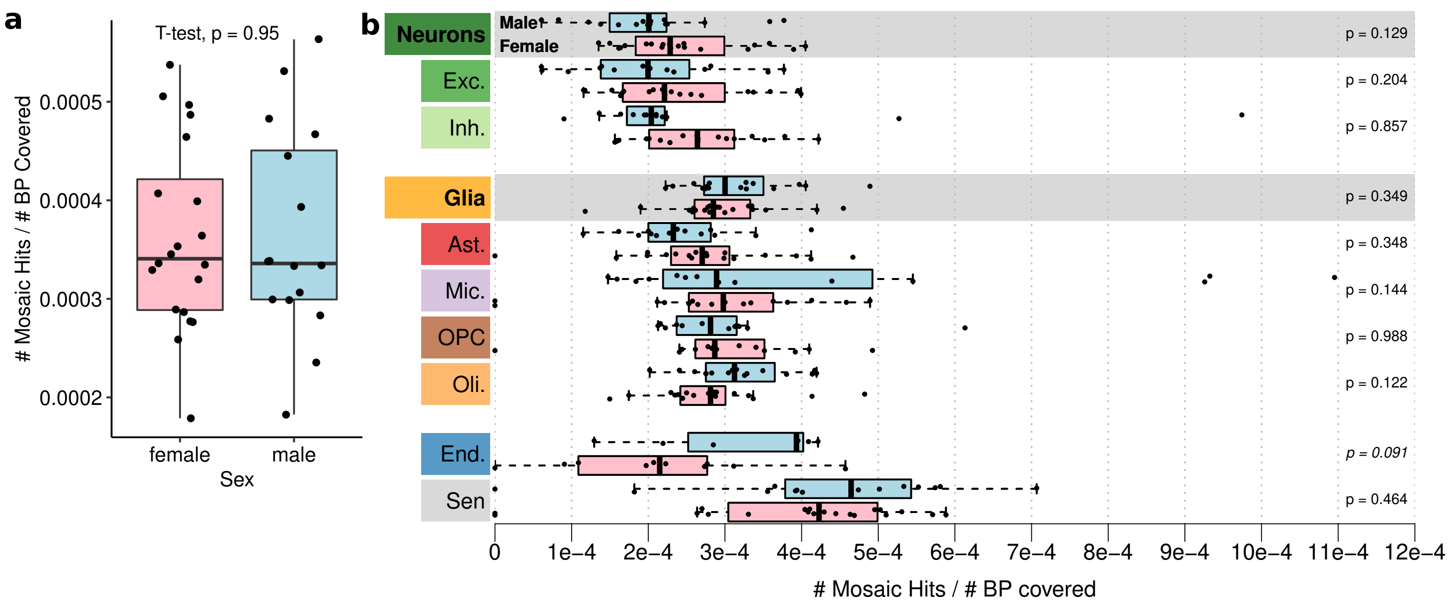


**Supplementary Figure 9. Mosaic mutational burden by sex. a.** Mosaic mutational burden (number of missense and loss-of-function mosaic hits over number of callable bases) by sex across all cells in 36 individuals. P-value is reported for two-sided t-test. **b.** Per-cell-type mosaic mutational burden by sex (female in pink, male in blue), for each cell type and neuronal and glial cells overall. P-values for two-sided t-tests are reported.

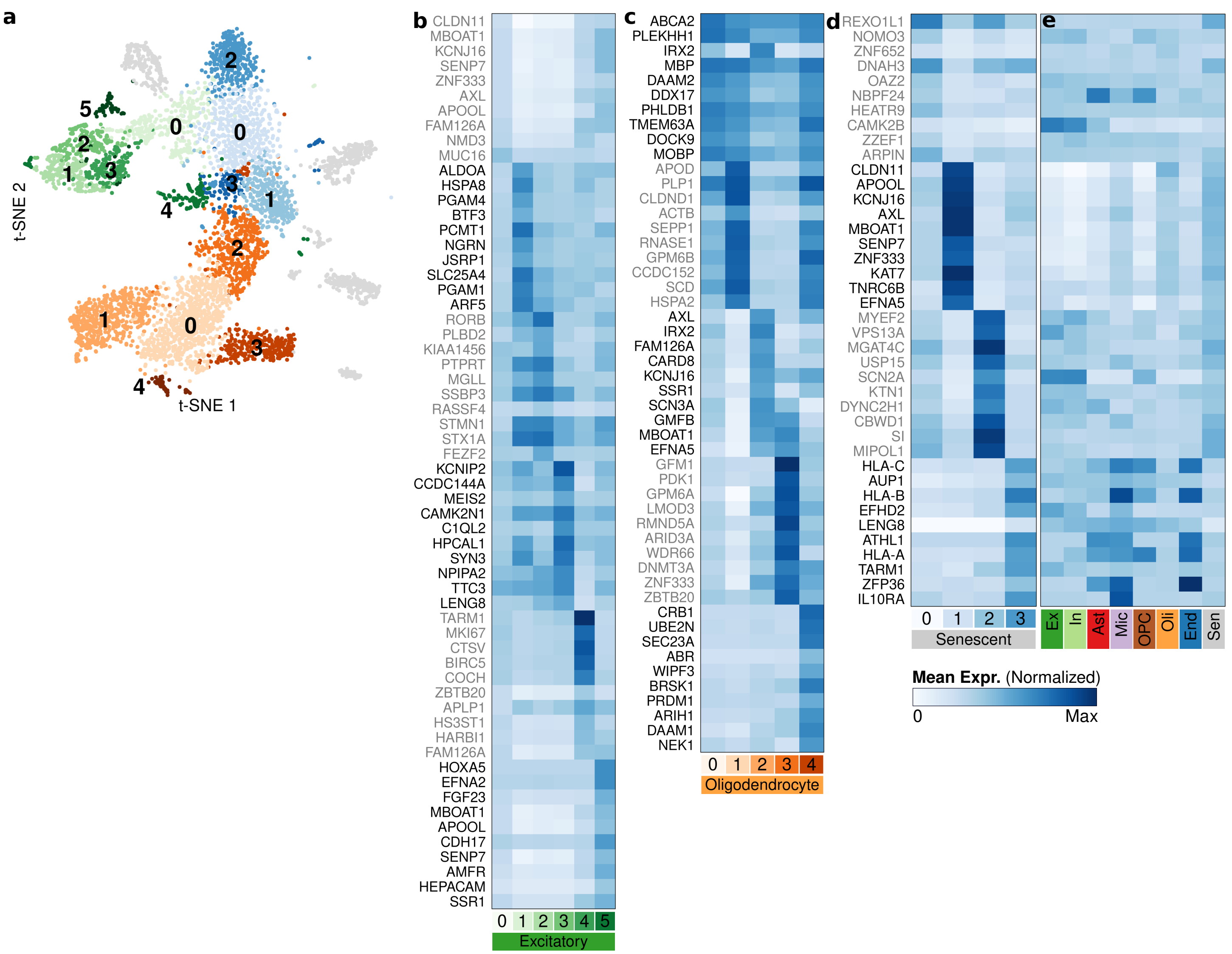

**Supplementary Figure 10. Major cell type subclusters and marker genes. a.** Subclusters for excitatory neurons, oligodendrocytes, and senescent cells. **b-d.** Average expression of the top 10 markers for each excitatory, oligodendrocyte, or senescent subcluster. Marker genes were computed for each subcluster against all cells in the main cluster by a wilcoxon test. **e.** Average expression of each of senescent subclusters' markers in the major cell types.

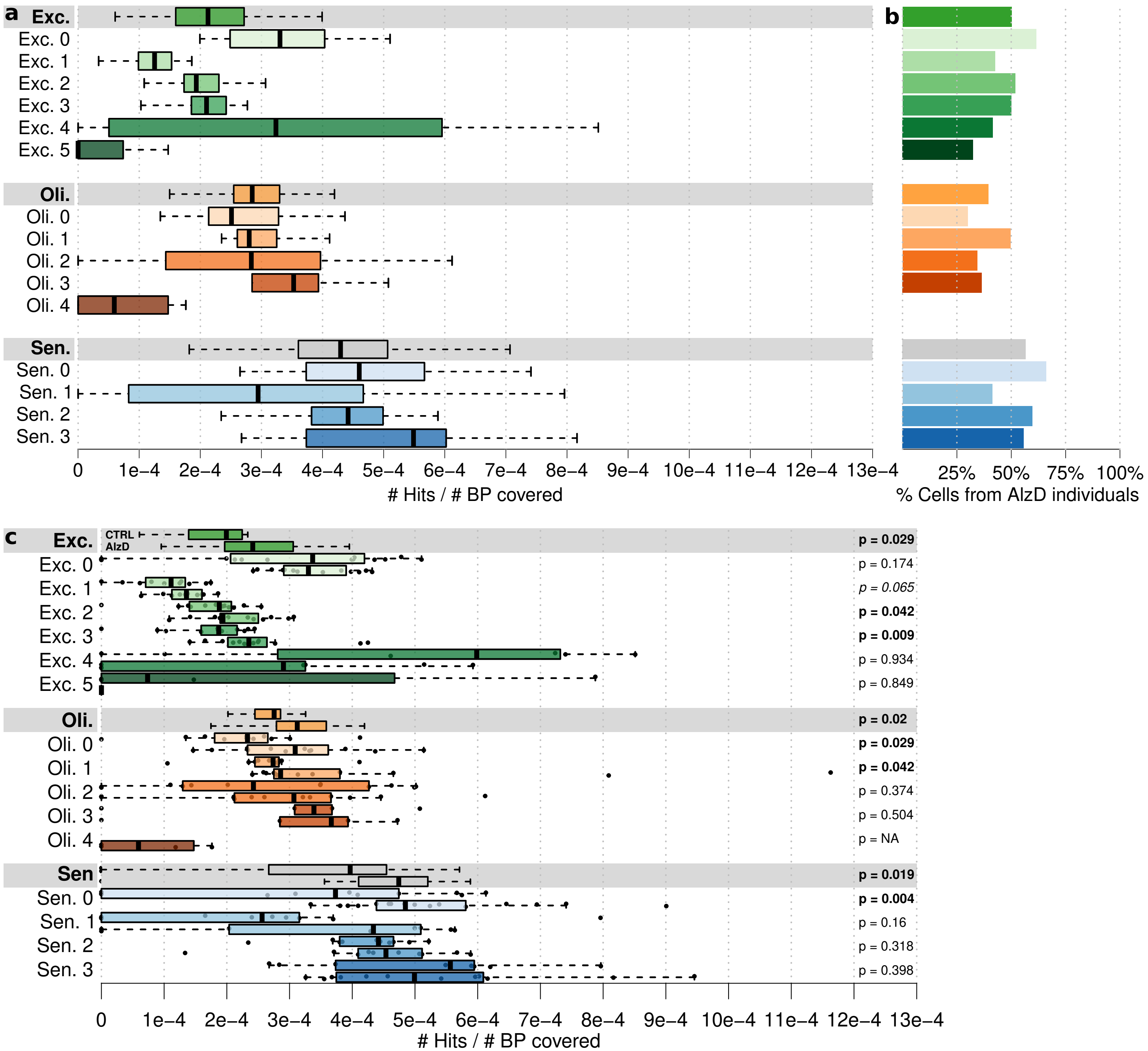


**Supplementary Figure 11. Mosaic mutational burden in subclusters in disease. a.** Mosaic mutational burden (number of missense and loss-of-function mosaic hits over number of bases at a callable depth) for all 36 individuals in each of excitatory neurons, oligodendrocytes, and senescent cells and their subclusters. **b.** Percentage of cells in each set from Alzheimer's dementia individuals. **c.** Mosaic mutational burden by subcluster and Alzheimer's dementia status for each of excitatory neurons, oligodendrocytes, and senescent cells and their subclusters.

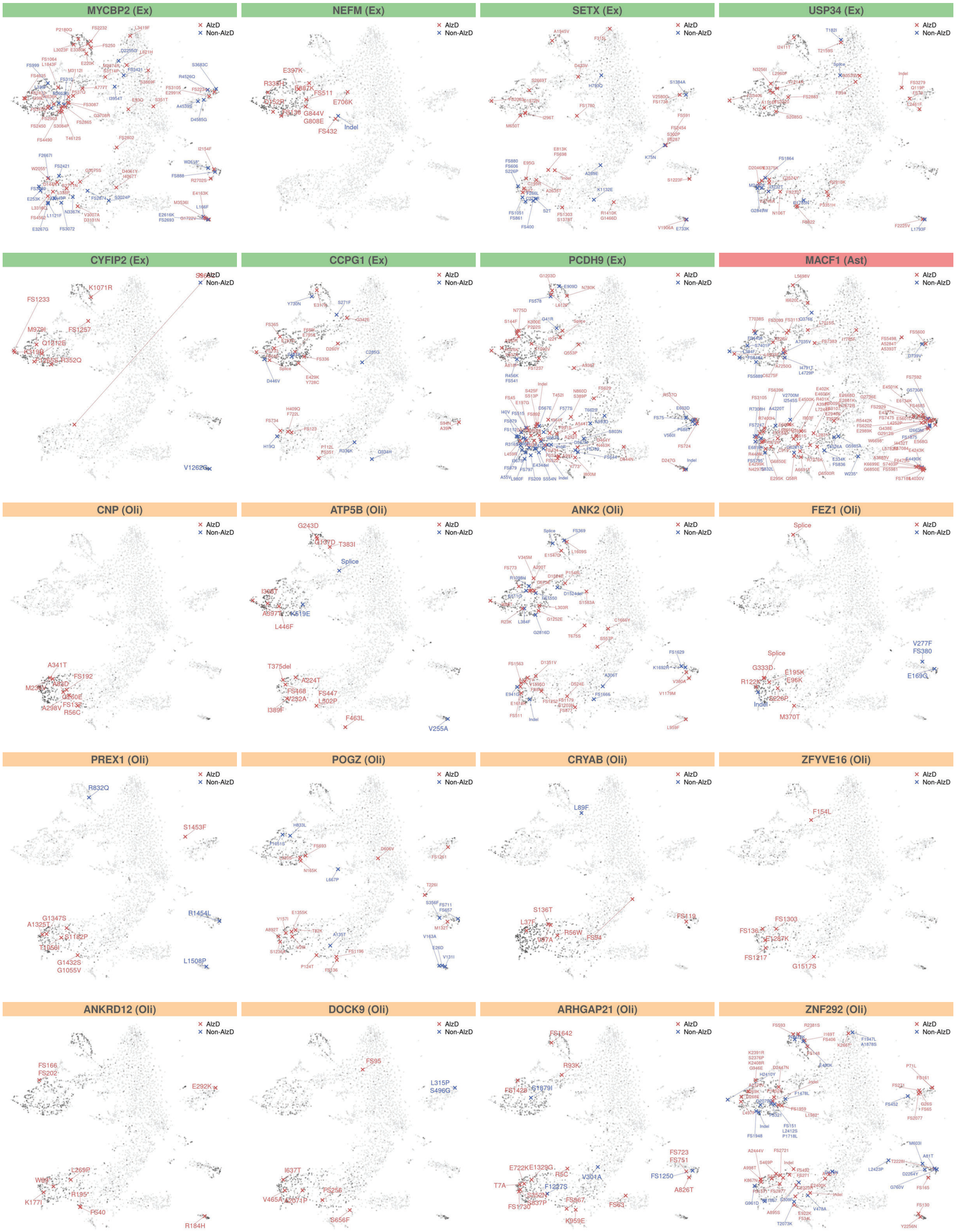


**Supplementary Figure 12. Genes with significantly higher burden in AlzD.** Mosaic hits in each over-burdened gene in excitatory neurons, astrocytes, and oligodendrocytes are labeled with their change and colored by their occurrence in cells from individuals with Alzheimer's dementia or no dementia. The t-SNE is colored according to coverage of the gene, with darker colors indicating higher coverage. All missense and LoF mutations used in burden calculations are shown, even in cell types where the gene is not enriched for burden.

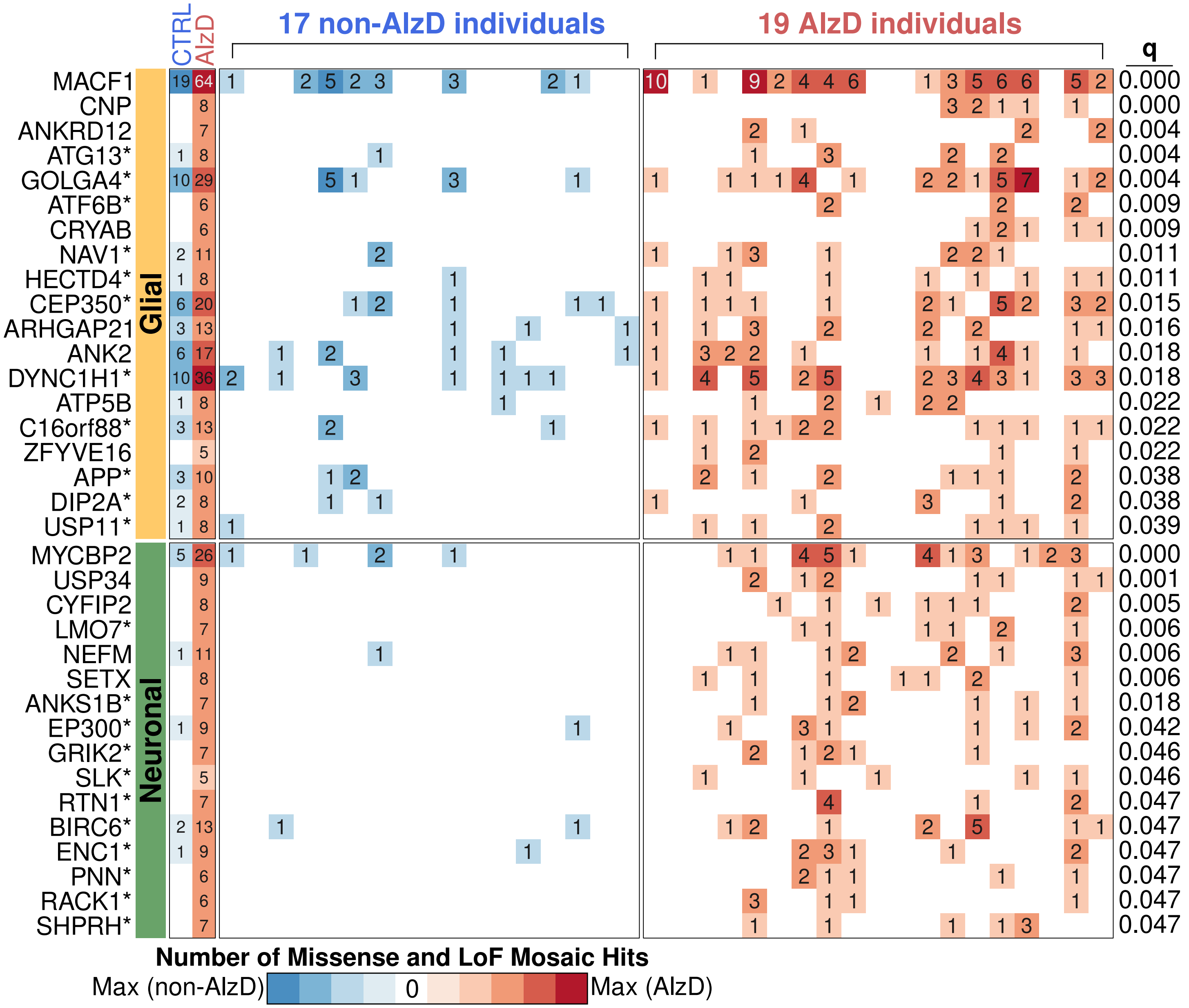


**Supplementary Figure 13. Neuronal and Glial genes enriched for AlzD burden.** Genes showing increased mosaic mutational burden (missense and loss-of-function hits, binomial test and FDR correction, q <0.05 shown) in Alzheimer's dementia versus non-dementia individuals in aggregated glial cells and neuronal cells. Heatmaps show number of mutations (left) overall in AlzD and non-AlzD individuals, (center) in each non-AlzD individual, and (right) in each AlzD individual in the cohort. Stars indicate genes not highlighted in a single-celltype analysis.

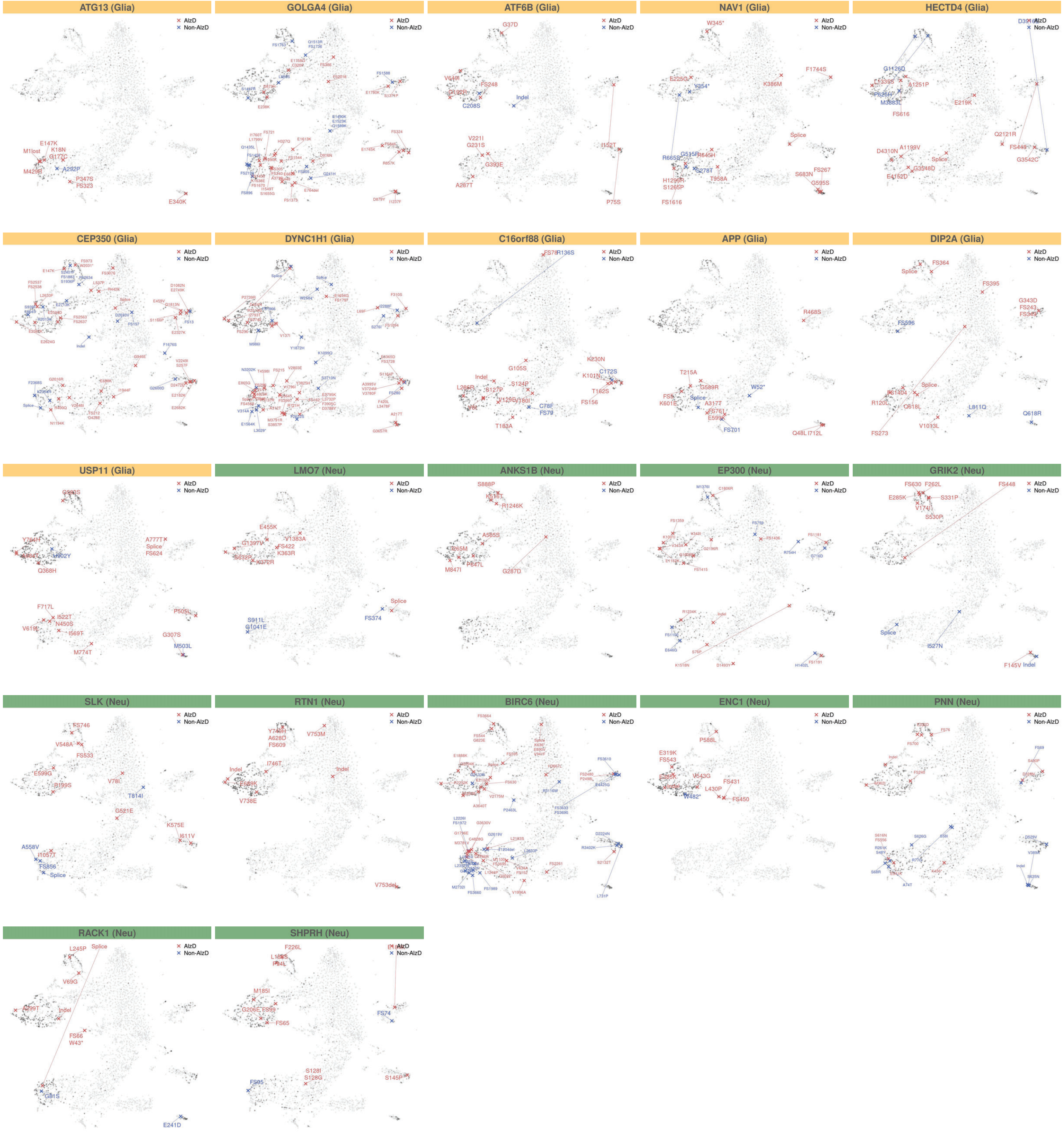


**Supplementary Figure 14. Genes with significantly higher burden in AlzD in neuronal and glial cells.** Mosaic hits in each over-burdened gene in neuronal and glial cells are labeled with their change and colored by their occurrence in cells from individuals with Alzheimer's dementia or no dementia. The t-SNE is colored according to coverage of the gene, with darker colors indicating higher coverage. All missense and LoF mutations used in burden calculations are shown, even in cell types where the gene is not enriched for burden.

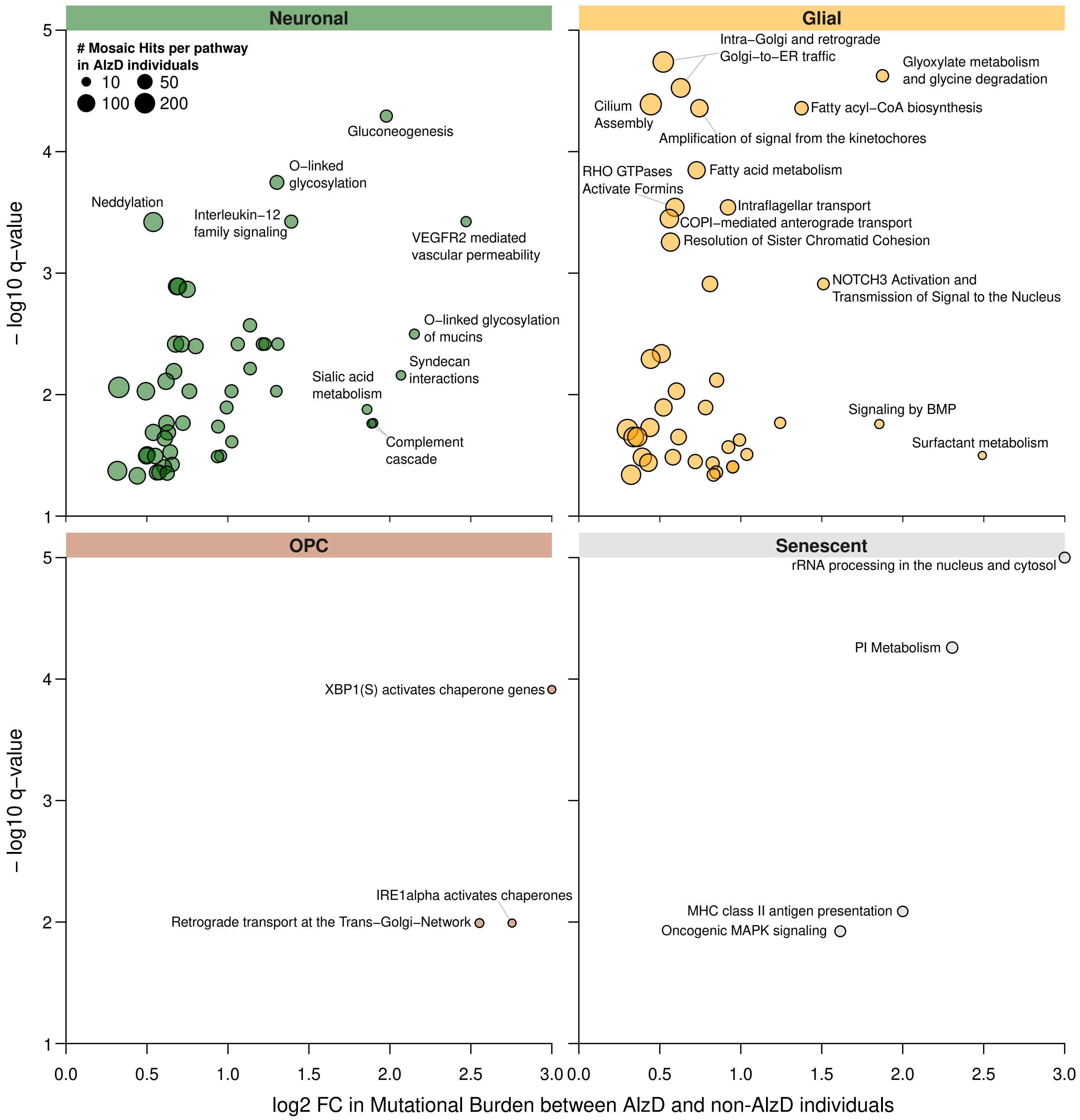


**Supplementary Figure 15. Pathway-level enrichments in disease.** Pathways with increased mosaic mutational burden in AlzD versus non-AlzD individuals in either neuronal cells, glial cells, OPCs, or senescent cells (missense and loss-of-function hits, log2 fold-change in burden against -log10q-value, size corresponds to number of hits in the pathway). Pathways with q <0.001 or log2fc >1.5 are labeled. Only pathways withstanding FDR correction with q <0.05 for a binomial test against burden in non-AlzD individuals are shown (see Methods).

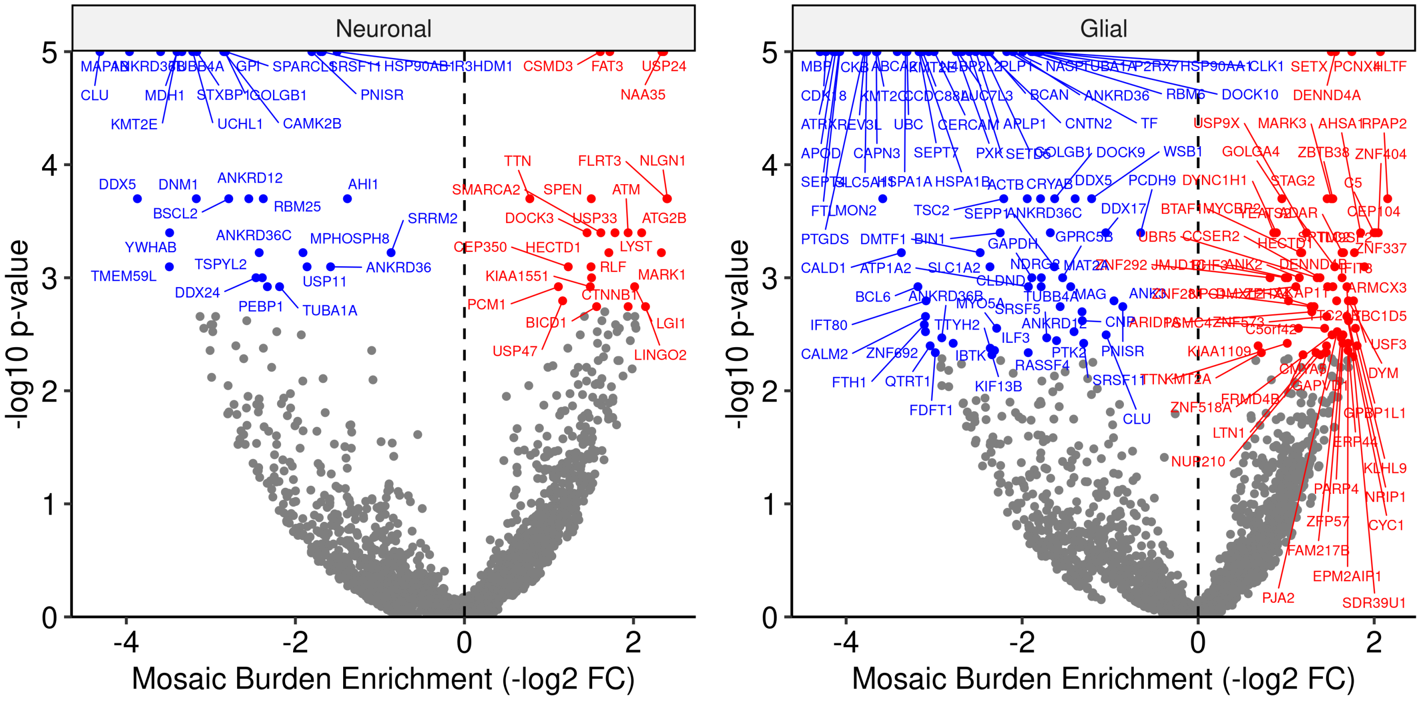


**Supplementary Figure 16.** Genes significantly depleted (blue) or enriched (red) for mosaic mutational burden in a disease-agnostic manner in neuronal cells or glial cells (missense and loss-of-function hits, log2 fold-change in burden against -log10p-value; genes with q <0.05 are labeled).

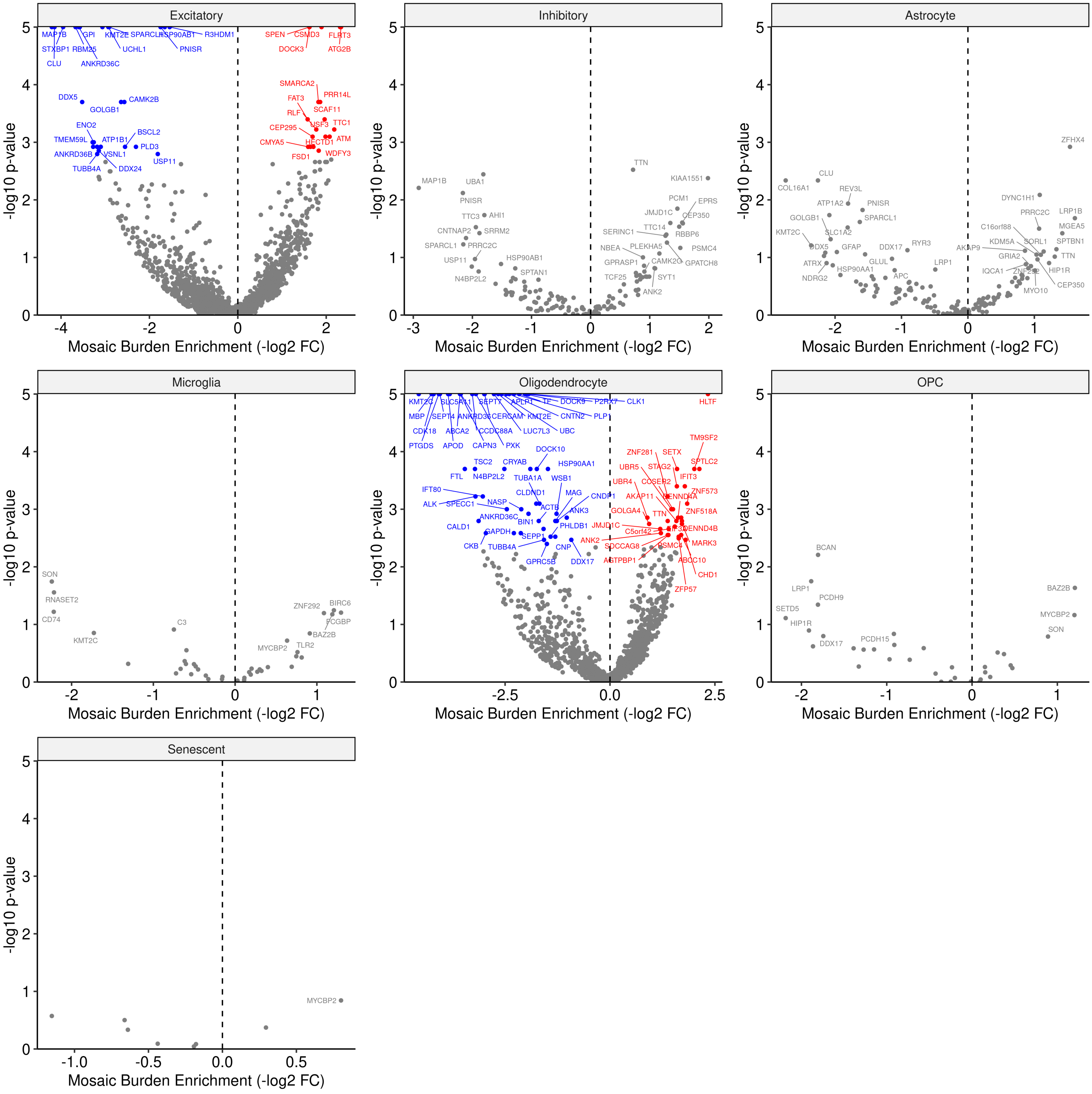


**Supplementary Figure 17.** Genes significantly depleted (blue) or enriched (red) for mosaic mutational burden in a disease-agnostic manner in each of the captured cell types (missense and loss-of-function hits, log2 fold-change in burden against -log10p-value; genes with q <0.05 are labeled in red/blue; genes with nominal p <0.1 are labeled in grey).
